# Multiomics reveals epigenetic control of fibroblast activity after myocardial infarction and a key role for RUNX transcription factors

**DOI:** 10.64898/2026.01.12.699153

**Authors:** Yuxia Li, Xujia Zhang, Kishan Ghimire, Nishan Khatri, Leshan Wang, Qianglin Liu, Mohammad Shiri, Rayyan Shakoori, Sachin Upadhayaya, Boyu Li, Joseph Francis, Constantine A. Simintiras, Rui Li, Jiangwen Sun, Xing Fu

## Abstract

**Background:** After myocardial infarction (MI), cardiac fibroblasts proliferate and undergo a sequential differentiation process. They first transition into cardiac myofibroblasts, a transient and highly contractile state, and ultimately into matrifibrocytes, a more stable state that partially resembles chondrocytes. These dynamic transitions are essential for infarct healing and scar formation. While insufficient fibroblast activation can compromise infarct integrity, excessive activation promotes pathological fibrosis that impairs cardiac function. Despite its clinical importance, the transcriptional and epigenetic regulation of these transitions remain poorly understood. Elucidating underlying mechanisms is critical for developing strategies to fine-tune fibroblast activity during cardiac repair.

**Methods:** We performed bulk RNAseq, ATACseq, CUT&Tag, CUT&RUN, EMseq, and Hi-C on cardiac fibroblasts from uninjured and post-MI mouse hearts. In parallel, we conducted single-nucleus multiomic (snRNAseq and snATACseq) profiling across multiple time points after MI. Subsequent integrated analysis explored epigenetic mechanisms regulating cardiac fibroblast gene expression and activity. Using an improved computational strategy, we constructed gene regulatory networks to identify key transcription factors and biological processes regulated by these transcription factors. To assess the role of *Runx1* specifically, we used tamoxifen-inducible, fibroblast-specific *Runx1* knockout mice to evaluate transcriptional, epigenetic, and functional outcomes with the same genomic tools and additional complementary assays.

**Results:** Cardiac fibroblasts undergo extensive chromatin remodeling after MI, which is highly correlated with changes in transcriptomic profiles. In contrast, the role of DNA methylation is relatively minor. Gene regulatory network analysis identified *Runx1* as a central regulator of cardiac fibroblast proliferation and matrifibrocyte differentiation. *In vitro* and *in vivo* validation confirmed *Runx1* as a key modulator of transcriptional and epigenetic changes in cardiac fibroblasts. *Runx1* KO reduced cardiac fibroblast proliferation, disrupted the myofibroblast-to-matrifibrocyte transition, and affected macrophage cytokine expression through altered cardiac fibroblast–macrophage communication. Fibroblast-specific *Runx1* knockout mice showed improved post-MI survival and reduced cardiac dilatation, especially in males. Simultaneous *Runx2* deletion further enhanced the effects of *Runx1* knockout.

**Conclusions:** Cardiac fibroblast activation and differentiation after MI are regulated by dynamic epigenetic changes. *Runx1* plays a pivotal role in modulating cardiac fibroblast activities, and its deletion improves cardiac repair by mitigating maladaptive fibroblast responses. By illuminating the centrality of *Runx1* in post-MI repair, this study identifies an actionable pathway for therapeutically steering fibroblast responses.

## INTRODUCTION

Cardiovascular diseases (CVDs) remain the leading cause of mortality worldwide, posing a major public health challenge. In 2017 alone, CVD-related healthcare costs and lost productivity in the US were estimated at $378 billion^1^. Among CVDs, acute myocardial infarction (MI) is particularly devastating due to its sudden onset, rapid progression, and poor prognosis^2^. MI is characterized by extensive cardiomyocyte (CM) death and subsequent tissue remodeling dominated by replacement fibrosis. While scar formation preserves structural integrity, persistent fibrosis impairs CM contractility, progressively reducing cardiac function and often culminating in heart failure^3–6^. A deeper understanding of this process is central to uncovering strategies to optimize fibrotic remodeling and improve post-MI outcomes.

Fibrosis in the heart is primarily driven by cardiac fibroblasts (CFs), a resident mesenchymal cell population with remarkable plasticity^7^. Our group and others demonstrated that following MI, CFs rapidly activate, proliferate, and differentiate into cardiac myofibroblasts (CMFs), which secrete large amounts of fibrillar collagen and acquire a smooth muscle-like phenotype through contractile protein expression^8–10^. Unlike regenerative tissues where myofibroblasts resolve via apoptosis or dedifferentiation, our previous work revealed that CFs persist within the infarct and transition into a previously unrecognized state termed matrifibrocytes (MaFs)^10–12^. MaFs exhibit reduced expression of CMF markers such as *Col1a1* and *Acta2* but upregulate cartilage and bone-associated genes, indicative of an advanced fibrotic stage. Beyond matrix deposition, CFs actively interact with immune cells and CMs, orchestrating cardiac remodeling after injury^13–15^.

Epigenetic pathways, such as DNA methylation, histone modification, transcription factor (TF), and chromatin interaction, tightly regulate gene expression. We recently showed that transcriptomic changes in CFs after MI are correlated with chromatin accessibility^16^. Studies from others also suggest an important role of epigenetics in CF activation and fibrosis^17–21^. However, systematic investigations and identification of novel regulators remain limited. Here, we integrate bulk and single-cell/nucleus transcriptomic and epigenomic approaches with fibroblast-specific lineage tracing to define a critical role for chromatin remodeling in this process. Our analysis highlights key regulators, including members of the runt-related transcription factor (Runx) family. Functional validation using models with fibroblast-specific deletion of Runx TFs further reveals their multifaceted roles and suggests potential therapeutic opportunities through targeted modulation of these factors.

## METHODS

### Animals

*Pdgfra^CreERT^*^2^ mice (JAX:032770)^22^ were crossed with *R26^GFP^* mice^23^ to generate *Pdgfra^CreERT^*^2^*;R26^GFP^* mice. To generate *Pdgfra^CreERT^*^2^*;Runx1^fl/fl^;R26^GFP^* and *Pdgfra^CreERT^*^2^*;Runx1^fl/fl^;Runx2^fl/fl^;R26^GFP^* mice, *Pdgfra^CreERT^*^2^*;R26^GFP^* mice were crossed with *Runx1^fl/fl^* (JAX:010673) or/and *Runx2^fl/fl^*^24^ mice. All mice are in C57BL/6J background. All experiments involving mice were approved by the Institutional Animal Care and Use Committee at Louisiana State University.

### Statistics

Unless otherwise specified, data are presented as mean ± SD with all data points shown in bar graphs or mean with whiskers showing Min to Max and all data points shown in box graphs. Statistical analyses were performed using Prism 10 (GraphPad Software, Inc.). For comparisons between two groups, a two-tailed t-test was applied. When comparing more than two groups, one-way ANOVA followed by Tukey’s post hoc test was used to assess statistical significance. Log-rank (Mantel-Cox) test was used to determine significance between the survival rates of *Pdgfrα^CreERT^*^2^*;R26^GFP^*, and *Pdgfrα^CreERT^*^2^*;Runx1^fl/fl^;R26^GFP^* mice. Significance levels are denoted as follows: *, p < 0.05; **, p < 0.01; ***, p < 0.001; ****, p < 0.0001.

### Data and code availability

Previous published bulk RNAseq and bulk ATACseq data are available at Gene Expression Omnibus (GEO: GSE186079). New raw bulk RNAseq, bulk ATACseq, bulk CUT&Tag, bulk CUT&RUN, bulk EMseq, and bulk Hi-C data have been deposited at GEO (GSE314445). All snMultiome and scRNAseq data have been deposited at GEO (SE315758). All original code for bulk sequencing data analysis has been deposited at GitHub (https://github.com/ODU-CSM/Pub-Runx1-25). All original data and code will be publicly available by the date of publication.

## RESULTS

### Bulk multiomic analysis reveals dynamic gene expression and epigenetic regulations in CFs after MI

To characterize transcriptomic changes in CFs after MI, we performed bulk RNAseq on lineage-traced CFs. The dataset combined previously published samples from uninjured hearts and hearts at 3 days (3D), 7 days (7D), 2 weeks (2W), and 4 weeks (4W) post-MI with newly collected samples at 2 hours (2H), 12 hours (12H), and 1 day (1D) post-MI, providing coverage across the entire remodeling timeline (**Figures 1A, S1A; Table S2**)^16^. Gene set enrichment analysis (GSEA) revealed early activation of RNA processing pathways as soon as 2H post-MI, peaking at 12H and persisting through 3D, indicating rapid transcriptional engagement (**Figure 1B**). Translational activation followed a similar but slightly delayed pattern, with peak activity at 1D post-MI (**Figure 1B**). Although CF proliferation becomes detectable at 3D post-MI, cell cycle-related pathways were activated by 2H, suggesting earlier CF activation than previously reported^10^. Actin organization terms remained enriched at 4W, despite declined expression of *Acta2*, the hallmark of CMFs, suggesting sustained cytoskeletal activity in MaFs (**Figures 1B, S1B**). Bone and cartilage-related pathways emerged by 7D, marking the onset of the transition from CMFs to MaFs (**Figure 1B**). Inflammatory and immune response pathways were also enriched, particularly at 7D, indicative of CF–immune cell interactions during remodeling (**Figure 1B**).

**Figure 1.**
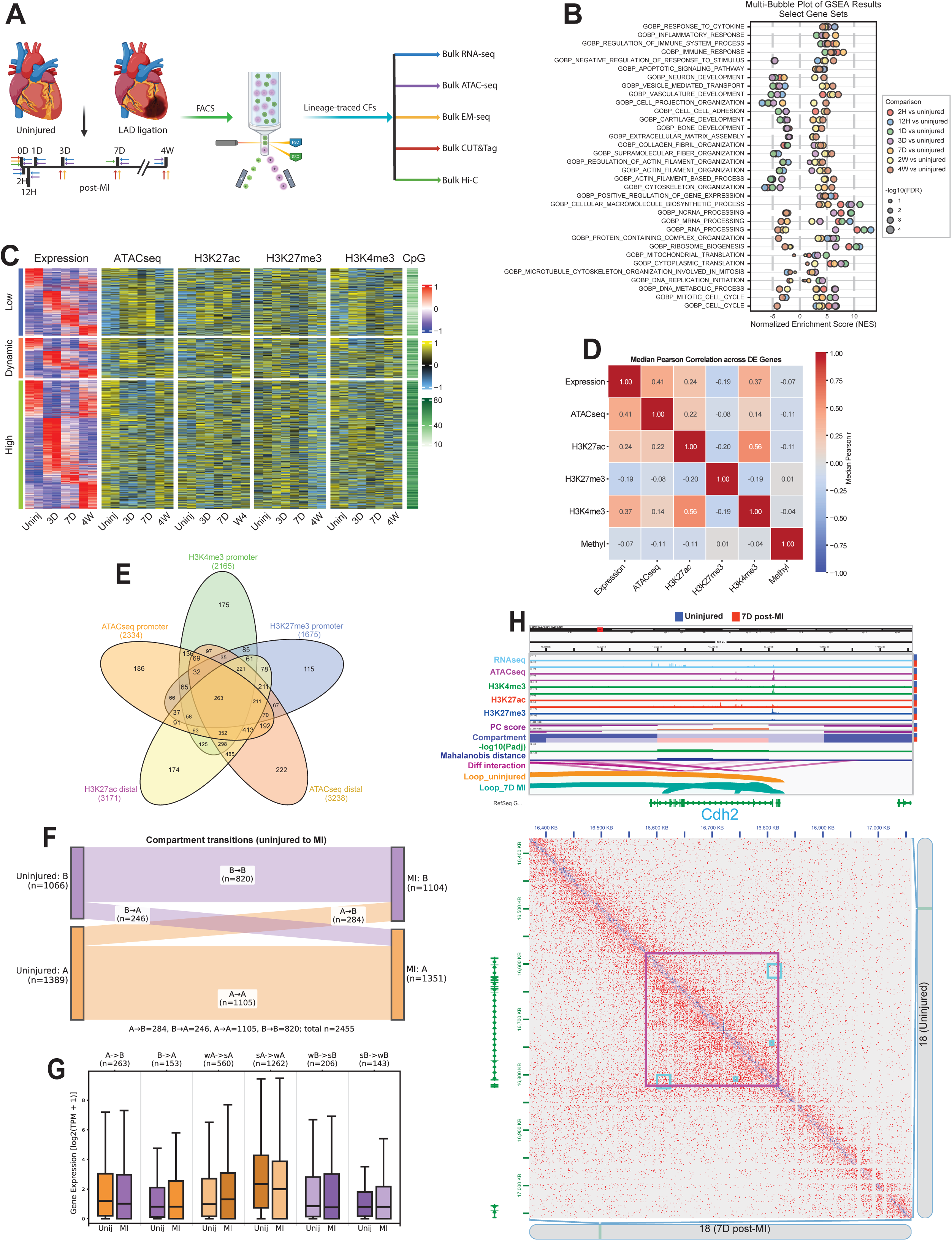
Bulk multiomic analysis reveals dynamic gene expression and epigenetic regulations in CFs after MI. (A) A schematic shows the experimental design in which CFs isolated from uninjured hearts and at different time points post-MI were subjected to the indicated genomic analyses. (B) A multi-bubble plot shows selected differentially enriched GOBP terms in indicated comparisons identified by GSEA. (C) A multi-modality heatmap shows the expression of DEGs across time points, along with promoter chromatin accessibility (ATACseq) and histone modification signals (H3K4me3, H3K27me3, H3K27ac). Genes are classified into high, low, and dynamic expression groups. Promoter CpG densities are also shown. (D) A heatmap shows the median Pearson correlations between gene expression and epigenetic marks in promoter regions of DEGs from uninjured, 3D, 7D, and 4W post-MI CFs. (E) A Venn diagram compares DEGs correlated with epigenetic marks (negative for H3K27me3, positive for others) in promoter or distal regions across uninjured and post-MI CFs. (F) A Sankey graph illustrates compartment changes between CFs from uninjured hearts and those at 7D post-MI identified by Hi-C. (G) A box plot shows the expression of genes associated with differential compartment transition. wA, weak A; sA, strong A; wB, weak B; sB, strong B. (H) An integrated visualization of CFs from uninjured and 7D post-MI hearts. The upper panel includes RNAseq, ATACseq, and histone Cut&Tag (H3K4me3, H3K27ac, H3K27me3) tracks, Hi-C tracks showing dcHiC-derived PC scores, compartments (dark red: sA; light red: wA; light blue: wB; dark blue: sB), compartment transition significance (-log10 Padj), Mahalanobis distance, and differential interactions, and Juicer-identified contacts. The lower panel shows Hi-C contact matrices for uninjured (above diagonal) and 7D post-MI (below diagonal) CFs, with purple boxes indicating TADs and cyan boxes indicating contacts exclusively identified at 7D post-MI.

To examine epigenetic regulation, we analyzed bulk ATACseq data from CFs at 2H, 12H, and 1D post-MI, along with previously published data from later time points (**Figure 1A, S1C; Table S3**)^16^. Approximately 24% of differentially expressed genes (DEGs) showed corresponding changes in promoter accessibility (**Figure S1D**), lower than the ∼40% reported when early time points were excluded^16^, suggesting additional regulatory mechanisms during initial activation. We next profiled histone modifications using CUT&Tag for H3K4me3 (active promoters), H3K27me3 (repressed promoters), and H3K27ac (active enhancers) in CFs from uninjured hearts and hearts at 3D, 7D, and 4W post-MI (**Figures S1E-G; Tables S4-6**). DEG expression correlated positively with promoter accessibility and H3K4me3 and H3K27ac signals and negatively with H3K27me3 (**Figures 1C-D, S1H**). Many distal regions with differential accessibility or H3K27ac enrichment, likely enhancers, also correlated strongly with nearby gene expression (**Figures S1I-L**). Most DEGs were associated with multiple chromatin marks, highlighting the complexity of CF gene regulation through coordinated chromatin remodeling (**Figure 1E**).

To assess DNA methylation, we performed enzymatic methyl-sequencing (EMseq) on CFs from uninjured hearts and hearts at 3D, 7D, and 4W post-MI (**Figure S1M**). Unlike H3K27me3, promoter methylation showed only a weak negative correlation with DEG expression across time points, suggesting a limited role in transcriptional regulation during CF differentiation (**Figures 1D, S1H**). In contrast, stronger negative correlations between DNA methylation and active chromatin marks than between H3K27me3 and active marks were identified across genes at individual time points, indicating that DNA methylation more broadly influences activity variability across loci (**Figure S1N**). No correlation was observed between H3K27me3 and DNA methylation, suggesting these repressive epigenetic mechanisms operate independently (**Figure 1D, S1N**).

Finally, to examine chromatin architecture, we performed Hi-C on CFs from uninjured hearts and hearts at 7D post-MI (**Figure 1A**). We identified around 2,500 differential compartment calls, including shifts between A and B compartments and sub-compartments (**Figure 1F, S1O**). These changes aligned with alterations in gene expression and other epigenetic marks, exemplified by the *Cdh2* locus, which encodes a cell adhesion protein^25^ (**Figure 1G-H**).

### Single-nucleus multiomic analysis reveals dynamic gene expression and chromatin accessibility in cardiac cell types after MI

To examine cardiac transcriptomic and chromatin accessibility changes after MI at single-cell resolution, we performed single-nucleus RNAseq and ATACseq (snMultiome) on uninjured hearts and hearts at 2H, 1D, 3D, 7D, and 4W post-MI (**Figure 2A**). Integrated analysis identified 19 clusters representing nine major cell types with dynamic abundance patterns, including CFs, cardiomyocytes (CMs), endothelial cells (EndoCs), mural cells (MuCs), monocytes/macrophages (Mo/Ma), endocardial cells (EndoCCs), epicardial cells (EpiCCs), T cells, and B cells (**Figures 2B-D**).

**Figure 2.**
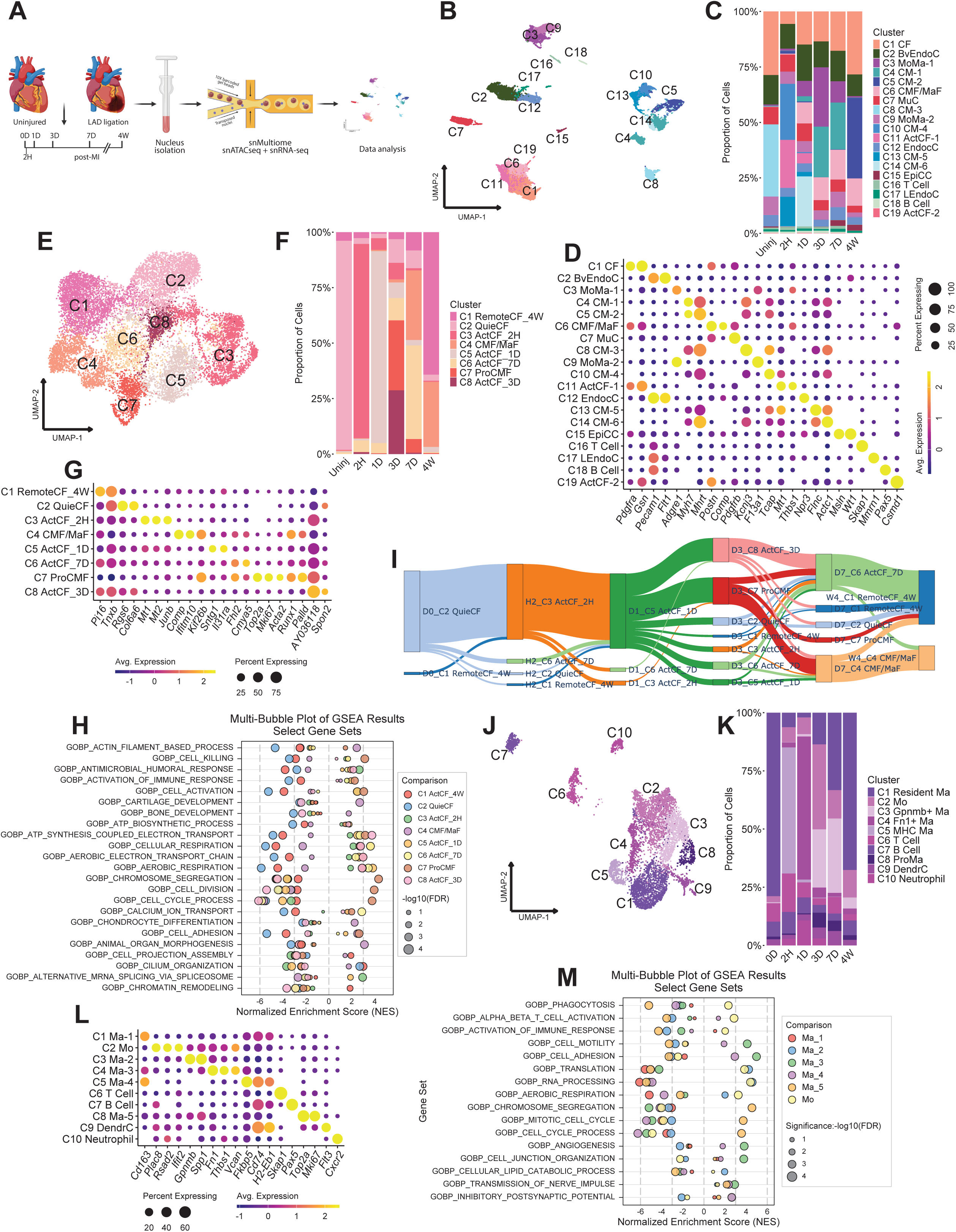
snMultiomic analysis reveals dynamic gene expression and chromatin accessibility in cardiac cell types after MI. (A) A schematic shows the experimental design in which nuclei isolated from uninjured hearts and at indicated time points post-MI were subjected to snMultiomic analysis. (B-C) A multi-modality joint UMAP graph (B) and a bar graph (C) show different cell types and subtypes identified in mouse hearts without injury and at different time points post-MI. (D) A dot plot displays the expression of representative marker genes used for cell type annotation. (E-F) A joint UMAP graph (E) and a bar graph (F) show CF subtypes identified across time points. (G) A dot plot shows the expression of representative marker genes used for CF subtype annotation. (H) A multi-bubble plot illustrates differential enrichment of selected GOBP terms in CF subtypes identified by GSEA. (I) CF differentiation trajectory generated using the MOSCOT pipeline. (J-K) A multi-modality joint UMAP graph (J) and a bar graph (K) show ImmunC types and subtypes identified across time points. (L) A dot plot displays expression of representative marker genes used for ImmunC annotation. (M) A multi-bubble plot shows differential enrichment of selected GOBP terms in Mo/Ma subtypes identified by GSEA.

CF reclustering revealed eight subclusters, including quiescent CFs (C2 QuieCF, *Rgs6*^+^), proliferating CMFs (C7 ProCMF, *Acta2*^+^,*Mki67*^+^), C4 CMF/MaF (*Palld*^+^,*Comp*^+^), remote CFs at 4W post-MI (C1 RemoteCF_4W, *Pi16*^+^), and four activated CF clusters enriched at specific time points (C3 ActCF_2H, *Junb*^+^; C5 ActCF_1D, *Il31ra*^+^; C6 ActCF_7D, *Cmya5*^+^; C8 ActCF_3D, *Spon2*^+^) (**Figures 2E-G, S2A-B; Tables S7-S8**). Notably, CMFs at 7D and MaFs at 4W grouped together and were distinct from ProCMFs at 3D (**Figures 2E-G**). *Acta2* expression in CMFs at 7D was similar to MaFs at 4W, despite sustained *Palld* expression, contrasting with bulk RNAseq, which showed sustained *Acta2* elevation at 7D (**Figures S2A-B, S1B**). This discrepancy likely reflects snMultiome capturing only newly synthesized nuclear transcripts, indicating reduced *Acta2* transcription and an ongoing CMF-to-MaF transition by 7D post-MI. ATACseq-inferred gene activity supported the reduced *Acta2* activity in CMF/MaF and identified elevated activity of additional MaF marker genes with low expression, such as *Chad* and *Cilp2* (**Figures S2C-D**). GSEA revealed activation of transcription, metabolism, and defense responses in early ActCF clusters (C3 and C5), proliferation signatures in ProCMFs (C7), and smooth muscle and chondrogenesis pathways in CMF/MaF clusters (C4), consistent with bulk RNAseq (**Figures 2H, S2E**). Trajectory analysis confirmed a progression from QuieCF (uninjured), through ActCF (2H, 1D) and ProCMF (3D), to CMF/MaF (7D, 4W) states (**Figure 2I**).

Immune cell reclustering identified T cells, B cells, dendritic cells, monocytes, and five macrophage subclusters (**Figures 2J-L**). GSEA revealed enrichment of angiogenesis, motility, and neurotransmission pathways in macrophage clusters C5 Ma-4 (*Fkbp5*^+^) and C4 Ma-3 (*Fn1*^+^), both *Cd163*^+^ resident macrophages abundant at 2H and 1D post-MI, suggesting roles in vascular and neuronal regulation early after MI (**Figures 2K-M**). Monocytes (C2 Mo, *Plac8*^+^) increased sharply at 3D, while monocyte-derived macrophages (C3 Ma-2, *Gpnmb*^+^) were enriched at 3D and 7D, showing strong activation of transcription, translation, and metabolism. Monocytes also displayed immune response and phagocytosis signatures, indicating active involvement in tissue repair during 3D-7D post-MI (**Figures 2K-M**). A proliferative macrophage population (C8 Ma-5) appeared primarily at 3D and 7D (**Figures 2K-M**). By 4W, most monocytes and monocyte-derived macrophages had receded, leaving *Cd163*^+^ resident macrophages as the dominant subtype (**Figure 2K**).

Dual-modality clustering identified nine CM clusters (**Figures S2F-H**). GSEA revealed rapid activation of immune-related pathways and actin remodeling in CMs as early as 2H post-MI (C3 CM-3), suggesting roles in immune signaling and early contractile adaptation (**Figure S2I**). Reclustering of mural cells yielded seven clusters, including one vascular smooth muscle cell (VSMC) cluster and six pericyte (PeriC) clusters, while EndoCs formed seven clusters consist of endocardial, lymphatic, and blood vessel subtypes (EndoCC, LvEndoC, and BvEndoC) (**Figures S2J-Q**). Similar to macrophages, proliferating clusters emerged in mural and endothelial cells at 3D and 7D and persisted at 4W, indicating a longer proliferative phase for vascular cells compared to CFs. Early metabolic adaptation, including activation of catabolic processes, was also evident in these cell types (**Figures S2J-Q**).

### Construction of gene regulatory networks regulating CF gene expression after MI

To identify TFs regulating CF gene expression after MI, we constructed a gene regulatory network (GRN) using CFs from the snMultiome dataset and the SCENIC+ pipeline^26^. The analysis was first performed on data normalized with the “sctransform” method, which improves sequencing depth correction^27^ (**Figures 3A-B, S3A**). However, because this approach compresses low-abundance signals toward zero, sensitivity for low-expression TFs and targets may be reduced. To address this, we repeated the analysis using standard log-normalization. Both methods identified many shared dynamic regulons across CF clusters, but regulons derived from log-normalized data were generally larger than those from sctransform-normalized data (**Figures 3A, S3B**). Among regulons with differential activity, Runx1+ (composed of genes positively regulated by *Runx1* and *Runx1* itself) showed strong and specific activation in C4 CMF/MaF and C7 ProCMF clusters (**Figures 3A, S3B**). This pattern, together with elevated *Runx1* expression in these clusters, suggests a key role for *Runx1* in CF differentiation after MI (**Figures S3C-D**).

**Figure 3.**
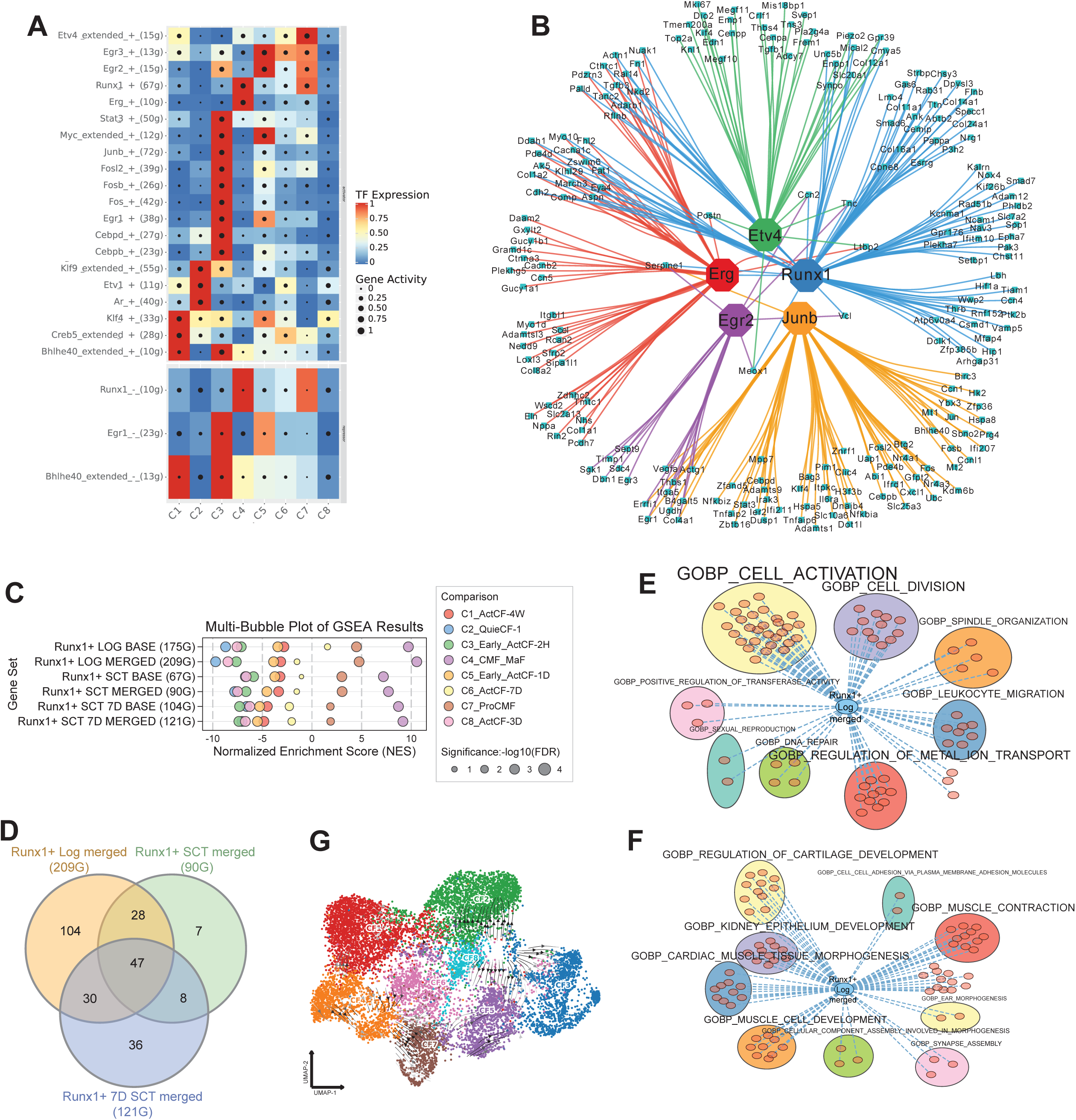
Construction of GRNs governing CF gene expression after MI. (A) A heatmap dot plot shows the gene-based regulon AUC scores and corresponding TF expression for the most differentially active regulons across CF subtypes identified using log-normalized data. (B) A Cytoscape network illustrates selected TFs and their target genes identified in the GRN. (C) A multi-bubble plot shows enrichment of different versions of Runx1+ regulon in CF clusters from uninjured and post-MI hearts. (D) A Venn diagram shows overlaps among indicated versions of the Runx1+ regulon. (E) A graph shows overlap between “Runx1+ Log merged” regulon and core enrichment genes in GOBP terms enriched in C7 ProCMF. (F) A graph shows overlap between “Runx1+ Log merged” regulon and core enrichment genes in GOBP terms enriched in C4 CMF/MaF. (G) A UMAP graph illustrates the effect of *Runx1* OE on CF differentiation trajectory predicted by SCENIC+.

A limitation of the standard SCENIC+ pipeline is its reliance on regulatory regions within ±150 kb of the transcription start site (TSS), which can miss distal enhancers. To address this, we incorporated topologically associated domains (TADs) identified by Hi-C, enabling detection of co-accessibility between promoters and enhancers located more than 150 kb apart but within the same TAD. In a second approach, we replaced snATACseq data with bulk H3K27ac CUT&Tag profiles to capture enhancer activity changes not reflected in chromatin accessibility. These modifications expanded the Runx1+ regulon to 209 genes in the log-normalized version (Runx1+ Log merged) and 90 genes in the sctransform-normalized version (Runx1+ SCT merged) (**Figures 3C, S3E; Table S9**). Analysis restricted to CFs at 7D post-MI identified 120 Runx1 targets (Runx1+ 7D SCT merged) (**Figures 3C, S3E; Table S9**). While substantial overlap was observed among the three versions, notable differences were present (**Figure 3D**). In particular, the divergence between “Runx1+ SCT merged” and “Runx1+ 7D SCT merged” suggests a differentiation state-specific role for *Runx1* in CFs. Moreover, GSEA showed stronger enrichment of “merged” version of Runx1+ regulons than the original version (“base”) in CMF/MaF, suggesting improved performance (**Figure 3C**).

Overlap analysis between Runx1 targets and core enrichment genes in Gene Ontology Biological Process (GOBP) terms enriched in ProCMFs indicated roles for *Runx1* in CF activation and proliferation in early post-MI stage (**Figures 3E, S3F**). Similar analysis for CMF/MaF suggested that *Runx1* promotes CMF and MaF differentiation at later stages, supported by enrichment of muscle contraction and cartilage development-related terms (**Figures 3F, S3G**). To assess the impact of *Runx1* modulation on CF gene expression, we simulated a *Runx1* overexpression scenario. This simulation shifted CFs toward CMF/MaF, the cluster with high Runx1+ regulon activity, reinforcing its role in guiding CF differentiation after MI (**Figures 3G, S3E**).

### *Runx1* promotes CF differentiation progress after MI

To further investigate the role of *Runx1* in CF activity after MI, we generated a tamoxifen-inducible, fibroblast-specific *Runx1* KO mouse line (*Pdgfra^CreERT^*^2^*;Runx1^fl/fl^;R26^GFP^*, abbreviated as *Runx1* KO). Single-cell RNA sequencing (scRNAseq) was performed on hearts of WT (*Pdgfra^CreERT^*^2^*;R26^GFP^*) and *Runx1* KO mice at 3D, 7D, and 4W post-MI (**Figure 4A**). Clustering analysis identified 15 clusters representing CFs, Mo/Ma, and other major cardiac cell types (**Figures 4B-C, S4A**).

**Figure 4.**
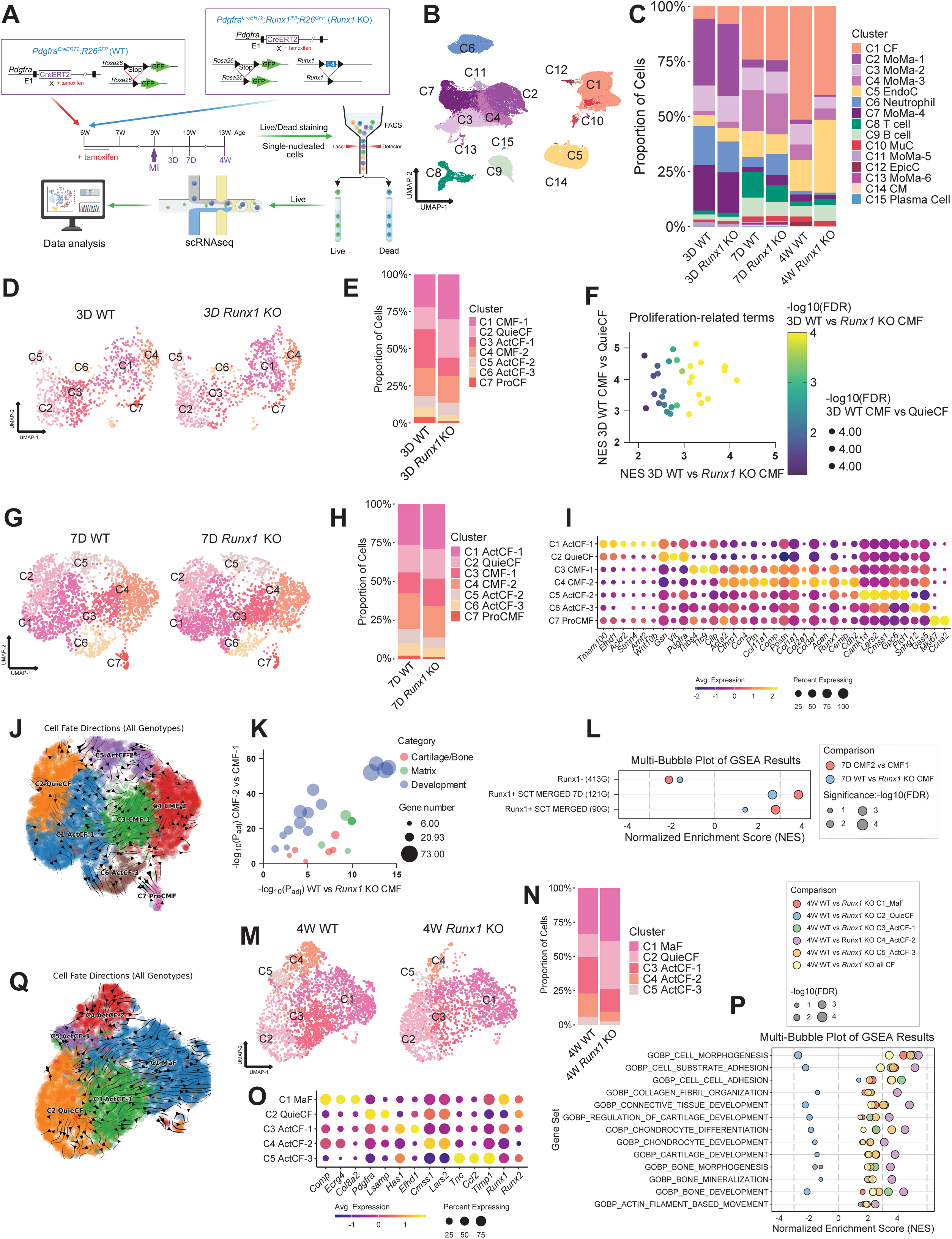
*Runx1* promotes CF differentiation after MI. (A) A schematic shows the experimental design in which viable cells isolated from WT and *Runx1* KO hearts at 3D, 7D, and 4W post-MI were subjected to scRNAseq and nuclei isolated from WT and *Runx1* KO hearts at 7D and 4W post-MI were subjected to snMultiome. (B) A UMAP graph (B) and a bar graph (C) show cell types and subtypes identified in WT and *Runx1* KO hearts at 3D, 7D, and 4W post-MI. (D-E) UMAP graphs (D) and a bar graph (E) show CF clusters identified in WT and *Runx1* KO hearts at 3D post-MI. (F) A multi-variable plot shows proliferation-related GOBP terms commonly enriched in WT versus *Runx1* KO CMFs and in CMFs versus QuieCFs at 3D post-MI. (G-H) UMAP graphs (G) and a bar graph (H) show CF clusters identified in WT and *Runx1* KO hearts at 7D post-MI. (I) A dot plot displays expression of selected genes in CF clusters at 7D post-MI. (J) A graph shows pseudotime trajectory of CFs at 7D post-MI. (K) A multi-variable plot highlights GOBP terms related to cartilage/bone formation, matrix, and development commonly enriched in CMF-2 versus CMF-1 and in WT versus *Runx1* KO CMFs at 7D post-MI. (L) A multi-bubble plot shows the differential enrichment of Runx1 regulons across indicated comparisons. (M-N) UMAP graphs (M) and a bar graph (N) show the CF clusters identified in WT and *Runx1* KO hearts at 4W post-MI. (O) A dot plot shows the expression of selected genes in CF clusters at 4W post-MI. (P) A multi-bubble plot shows differential enrichment of selected GOBP terms between WT and *Runx1* KO CFs of individual subtypes at 4W post-MI. (Q) A graph shows pseudotime trajectory of WT and *Runx1* KO CFs at 4W post-MI.

Reclustering CFs at 3D revealed an increase in QuieCF (C2) and a reduction in ActCFs (C3 ActCF-1, *Postn*^+^) and proliferating (C7 ProCF, *Mki67*^+^) CFs in *Runx1* KO hearts, consistent with the predicted role of *Runx1* in promoting CF activation and proliferation (**Figures 4D-E, S4B; Table S10**). When clustering was simplified to separate CMFs from QuieCFs, GSEA showed strong enrichment of proliferation-related terms in WT CMFs compared to *Runx1* KO CMFs, many of which were also enriched in CMFs versus QuieCFs at 3D (**Figures 4F, S4C**). Runx1+ regulon activity was significantly higher in WT ProCFs, and its overlap with core enrichment genes driving the enrichment of cell division-related terms in WT versus *Runx1* KO CMFs further supports the pro-proliferative function of *Runx1* (**Figures S4D-E**).

At 7D post-MI, CF reclustering identified seven clusters, including two CMF subtypes: C3 CMF-1 and C4 CMF-2 (**Figures 4G-I; Table S11**). *Runx1* KO hearts showed a lower CMF-2/CMF-1 ratio compared to WT hearts (**Figure 4H**). CMF-2 expressed higher levels of Runx1-targted CMF markers, such as *Acta2*, *Cemip*, and *Cdh2*, along with MaF-associated chondrogenesis genes (e.g., *Acan* and *Col11a1*), indicating more advanced differentiation than CMF-1 (**Figure 4I; Table S9**). Pseudotime analysis using CellRank2^28^ confirmed CMF-2 as a later stage in the differentiation trajectory (**Figure 4J**). GSEA revealed enrichment of cartilage/bone development and extracellular matrix terms in CMF-2 versus CMF-1, overlapping with terms enriched in WT versus *Runx1* KO CMFs (**Figure 4K**). Runx1+ regulons were enriched in CMF-2 and WT CMFs, while genes negatively regulated by Runx1 (Runx1-regulon), identified from a GRN constructed using bulk RNAseq and ATACseq data, were enriched in CMF-1 and *Runx1* KO CMFs (**Figure 4L**). These findings indicate that *Runx1* promotes CF progression toward differentiated states.

At 4W post-MI, CF reclustering identified five clusters, including C1 MaF, C2 QuieCF, and three ActCF clusters (C3–C5) (**Figures 4M-O; Table S12**). MaF abundance was similar between WT and *Runx1* KO hearts, but WT hearts contained more ActCFs, whereas *Runx1* KO hearts had more QuieCFs. This suggests that in chronic MI, *Runx1* KO does not significantly affect MaFs in the infarct core but may reduce CF activation in surrounding regions. Consistently, GSEA showed the greatest differences in MaF-related terms between WT and *Runx1* KO in C4 ActCF-2, the cluster with the largest abundance difference between WT and *Runx1* KO and second-highest expression of *Comp*, a chondrogenic gene^29^ targeted by *Runx1*, after C1 MaF (**Figures 4M-P, S4F; Table S9**). Pseudotime analysis positioned C4 ActCF-2 as a state parallel to C1 MaF, indicating its activated status (**Figure 4Q**). Overlap analysis between Runx1+ regulons and core enrichment genes in terms enriched in WT versus *Runx1* KO C4 ActCF-2 suggests that *Runx1* regulates pathways related to MaF differentiation, cell adhesion, development, and protein modification in CFs at 4W post-MI (**Figure S4G**). Trajectory analysis of CFs across all three post-MI time points using MOSCOT^30^ revealed distinct patterns between WT and *Runx1* KO hearts. Most WT CFs in “7D C1 ActCF-1” progressed to “4W C3 ActCF-1,” whereas a substantial fraction of *Runx1* KO CFs in “7D C1 ActCF-1” reverted to “4W C2 QuieCF” (**Figures S4H-I**). These findings suggest that in chronic MI, *Runx1* KO exerts a stronger effect on CFs exposed to submaximal activation stimuli.

### CF-specific *Runx1* KO ameliorates post-MI cardiac dysfunction

To assess the functional impact of CF-specific *Runx1* KO on post-MI healing, WT and *Runx1* KO mice were subjected to MI injury. *Runx1* KO significantly reduced rupture rates in male mice (**Figure 5A**). No difference was observed in female mice possibly due to their extremely low rupture incidence (**Figure S5A**). Echocardiography at 7D post-MI showed improved cardiac function and reduced ventricular dilation in male but not female *Runx1* KO mice compared to WT, suggesting a sex-dependent functional benefit (**Figures 5B-E, S5B-E**). Interestingly, infarct size was reduced in both male and female *Runx1* KO mice, suggesting a non-sex-specific antifibrotic effect (**Figures 5F, S5F**). By 4W post-MI, functional improvement was no longer evident, indicating a time-limited effect of *Runx1* KO (**Figures 5B-E, S5B-E**).

**Figure 5.**
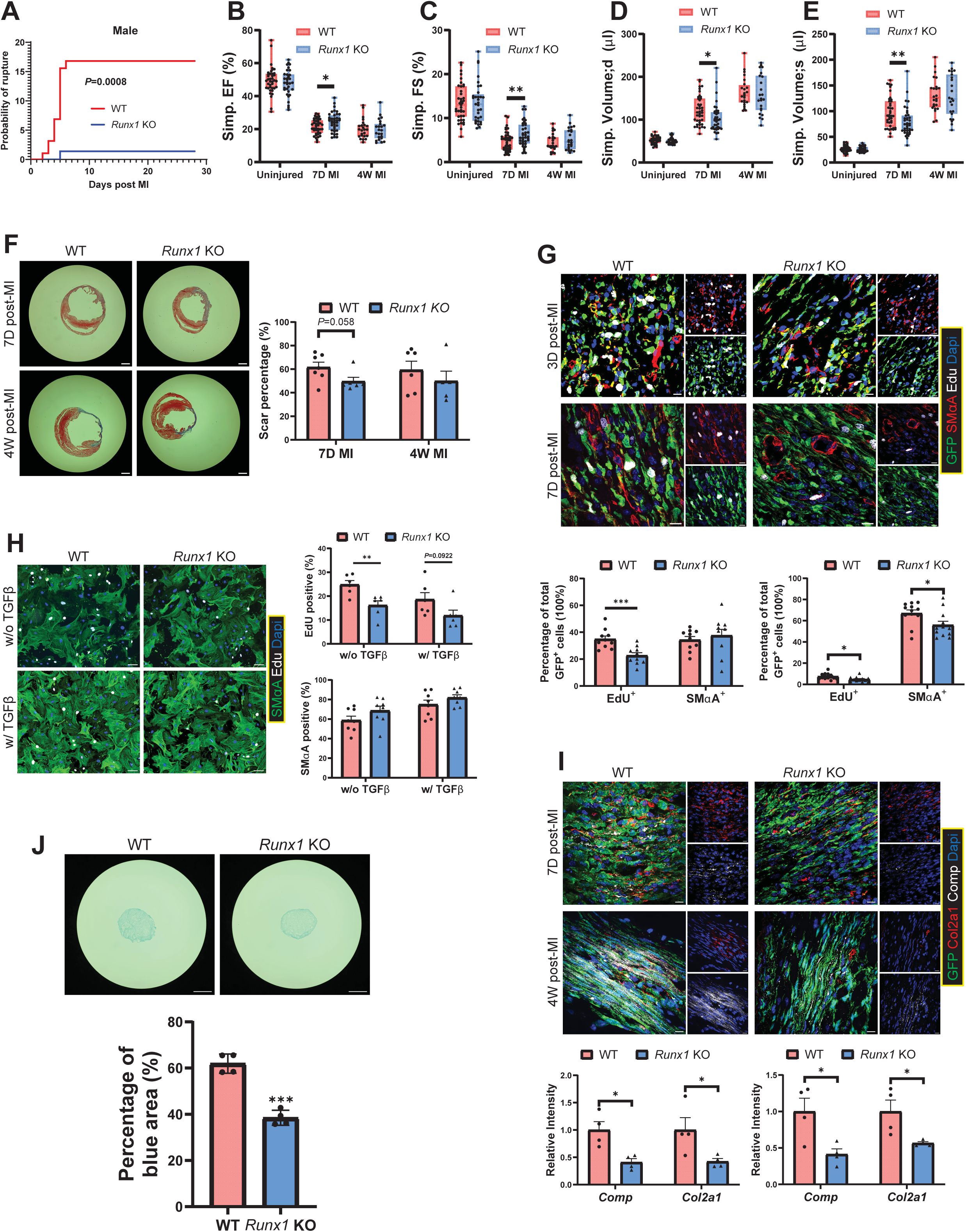
CF-specific *Runx1* KO ameliorates post-MI cardiac dysfunction. (A) Post-MI rupture curves for male WT and *Runx1* KO mice. n=95 (WT), 94 (*Runx1* KO). (B-E) Ejection fraction (B), fraction shortening (C), and diastolic (D) and systolic (E) volumes in male WT and *Runx1* KO mice measured at baseline and at 7D and 4W post-MI. n=39 (uninjured WT), 33 (uninjured *Runx1* KO), 41 (7D MI WT), 39 (7D MI *Runx1* KO), 24 (4W MI WT), 23 (4W MI *Runx1* KO). *, *p* < 0.05; **, *p* < 0.01. (F) Trichrome staining images show the infarcted hearts from male WT and *Runx1* KO mice at 7D and 4W post-MI with quantification of infarct size. Scale bar: 1 mm. n=7 (WT, 7D post-MI), 6 (*Runx1* KO, 7D post-MI), 6 (WT, 4W post-MI), 5 (*Runx1,* KO 4W post-MI). (G) IHC images of infarct regions at 3D and 7D post-MI stained for EdU and SMαA with quantification. Scale bar: 10 μm. *, *p* < 0.05; ***, *p* < 0.001. n=10 (WT, 3D MI), 10 (*Runx1* KO, 3D MI), 12 (WT, 4W MI), 12 (*Runx1* KO, 4W MI). (H) ICC images of cultured WT and *Runx1* KO CMFs stained for EdU and SMαA. Scale bar: 70 μm. **, *p* < 0.01. n=6 (EdU), 8 (SMαA). (I) IHC images of infarct regions at 7D and 4W post-MI stained for Comp and Col2a1 with quantification. Scale bar: 10 μm. *, *p* < 0.05. n=4. (J) Alcian blue staining of WT and *Runx1* KO CFs induced for chondrogenesis with quantification. Scale bar: 200 μm. *, *p* < 0.05. n=4.

Immunohistochemical staining (IHC) of 3D post-MI hearts and immunocytochemical staining (ICC) of cultured CMFs treated with or without TGFβ revealed reduced proliferation in *Runx1* KO CMFs compared to WT (**Figures 5G-H**), consistent with the pro-proliferative role of *Runx1* identified by sequencing analysis. Reduced smooth muscle alpha-actin (SMαA) expression in *Runx1* KO CMFs was observed at 7D post-MI (**Figure 5G**), which agreed with the Runx1 target identity of *Acta2* (**Table S9**). However, no difference was identified at 3D post-MI or *in vitro*, suggesting the presence of other factors regulating *Acta2* expression (**Figures 5G-H**). Immunostaining of 7D and 4W post-MI hearts showed reduced signals for chondrogenesis and MaF markers, Comp and Col2a1, both Runx1 targets, in *Runx1* KO hearts compared to WT, supporting a role for *Runx1* in MaF differentiation (**Figure 5I; Table S9**). Alcian blue staining of cultured CFs further confirmed attenuated chondrogenic activity in *Runx1* KO CFs compared to WT (**Figure 5J**).

### Multiomic analysis identifies epigenetic regulation of CF transcriptome by *Runx1*

To investigate how *Runx1* regulates CF transcriptional programs, we performed bulk multiomic analysis on cultured WT and *Runx1* KO CMFs, including RNAseq, ATACseq, and CUT&Tag profiling for H3K4me3, H3K27me3, and H3K27ac. RNAseq revealed 351 DEGs, including many Runx1 target genes, such as *Lox*, *Cemip*, *Col11a1*, *Col1a1* and *Ptn* (**Figure 6A; Table S9, S13**). Additional Runx1 targets also showed corresponding expression changes that did not reach the fold-change threshold (**Table S13**). Consistently, differential Runx1 regulon enrichment was pronounced between WT and *Runx1* KO CMFs (**Figure S6A**). GSEA identified clusters of GOBP terms enriched in WT versus KO CMFs, including those related to tissue remodeling, cell morphogenesis, immune response, and connective tissue development (**Figure S6B**). Overlap analysis between core enrichment genes led to the differential enrichment of these terms in WT CMFs and Runx1+ regulon suggests that *Runx1* directly promotes proliferation, MaF differentiation, stress response, and cell adhesion (**Figure 6B**), consistent with scRNAseq data.

**Figure 6.**
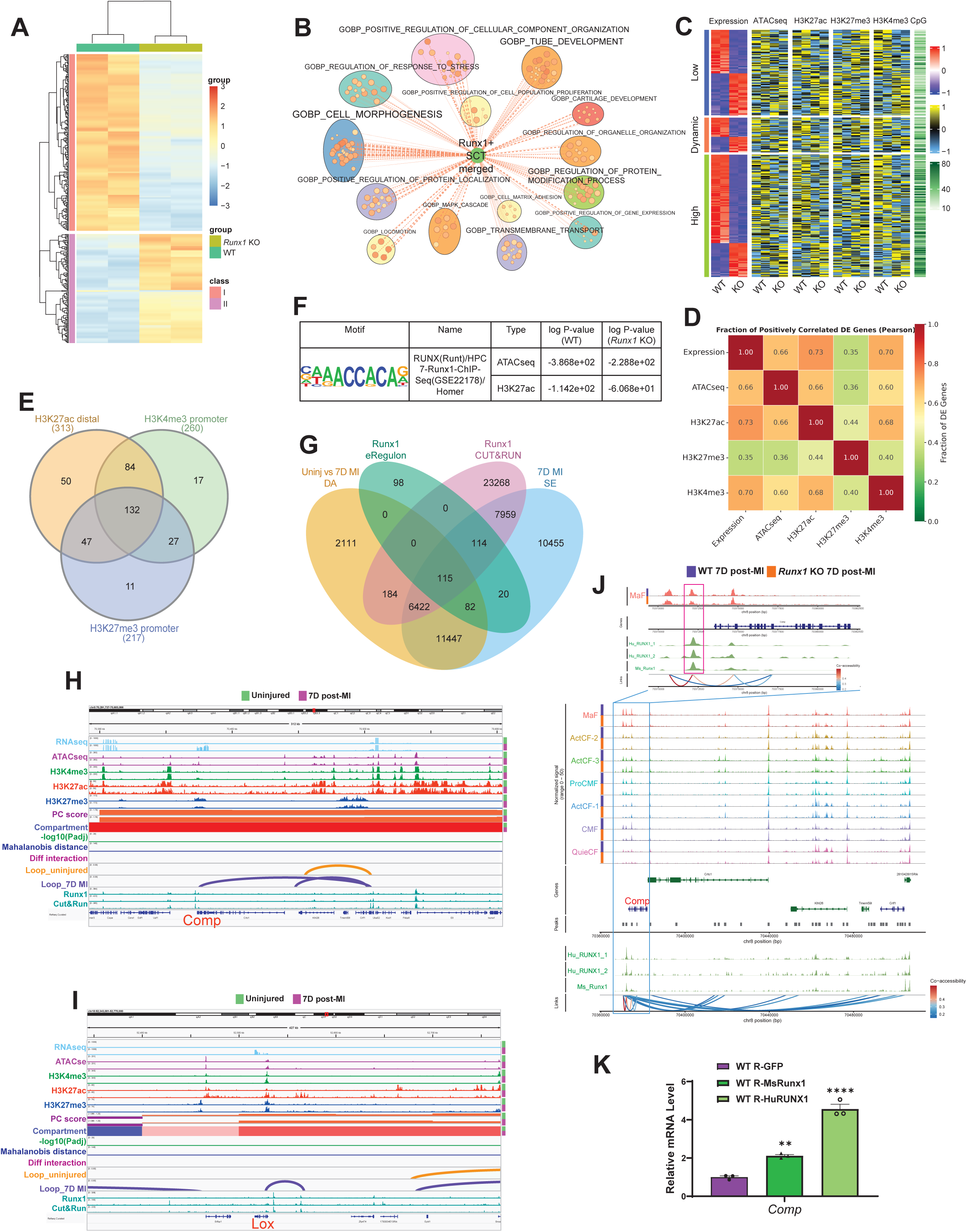
**Multiomic analysis identifies epigenetic regulation of CF transcriptome by *Runx1*.** (A) A heatmap shows differential gene expression between WT and *Runx1* KO CMFs. (B) A graph displays overlap between Runx1 regulon (Runx1+ SCT merged) and core enrichment genes in GOBP terms enriched in cultured WT versus *Runx1* KO CMFs. (C) A multi-modality heatmap shows the expression of genes differentially expressed between cultured WT and *Runx1* KO CMFs, and corresponding promoter chromatin accessibility (ATACseq) and histone marks (H3K4me3, H3K27ac, H3K27me3). Genes are classified into high, low, and dynamic expression groups. Promoter CpG densities are also shown. (D) A heatmap shows the fraction of DEGs between WT and *Runx1* KO CMFs with positive Pearson correlations between gene expression and epigenetic markers in promoter regions. (E) A Venn diagram shows overlap among WT versus *Runx1* KO DEGs positively correlated promoter H3K4me3 signal, negatively correlated with promoter H3K27me3 signal, and positively correlated with distal H3K27ac signal. (F) Runx1 motif enrichment P-values in distal ATACseq and H3K27ac CUT&Tag peaks of cultured WT and *Runx1* KO CMF. (G) A Venn diagram shows overlap among differential ATACseq peaks between uninjured and 7D post-MI CFs (Uninj vs 7D MI DA), Runx1-targeted enhancers identified by SCENIC+ (Runx1 eRegulon), Runx1 CUT&RUN peaks, and super enhancers identified in CFs at 7D post-MI (7D MI SE). (H-I) Integrated views for QuieCFs (uninjured) and CMFs (7D post-MI) show RNAseq, ATACseq, and histone CUT&Tag (H3K4me3, H3K27ac, H3K27me3) tracks, Hi-C tracks with dcHiC-derived PC scores, compartments (dark red: sA; light red: wA; light blue: wB; dark blue: sB), compartment transition significance (-log10 Padj), Mahalanobis distance, differential interactions, Juicer-identified contacts, and Runx1/RUNX1 CUT&RUN tracks. Panels show *Comp* (H) and *Lox* (I) loci. (J) A graph illustrates chromatin co-accessibility around the *Comp* locus based on snMultiome analysis of WT and *Runx1* KO CFs at 7D post-MI. The inset highlights a Runx1-targeted proximal enhancer located upstream of the *Comp* gene. (K) A bar graph shows the effects of overexpressing mouse *Runx1* (MsRunx1) and human *RUNX1* (HuRUNX1) OE on the expression of *Comp* in CFs.

Integrated transcriptomic and epigenetic analyses revealed coordinated changes in promoter accessibility and histone modifications for many DEGs, such as *Cemip*, between WT and *Runx1* KO CMFs (**Figures 6C-E, S6C-D; Table S14**). Motif analysis of ATACseq and H3K27ac CUT&Tag data showed a marked reduction in Runx1 motif enrichment within distal regulatory regions, primarily enhancers, in KO CMFs compared to WT (**Figure 6F**). To further examine enhancer targeting, we performed Runx1 CUT&RUN. Overlap analysis indicated that 229 of 429 Runx1-targeted enhancers predicted by SCENIC+ were directly bound by Runx1 (**Figure 6G**). Many of these enhancers were also located within CMF super-enhancers and/or overlapped regions with differential accessibility between QuieCFs and CMFs, indicating the critical role of *Runx1* in CF gene regulation and activation (**Figure 6G**). Integration with Hi-C data revealed increased Runx1-mediated enhancer–promoter loops and chromatin interactions in CMFs compared to QuieCFs for fibrosis-related Runx1 targets, such as *Comp*, *Lox*, *Cemip*, and *Cdh2*, most of which exhibited reduced expression in *Runx1* KO CMFs (**Figures 6H-I, S6E-G; Tables S9, S13, S15**). Consistently, snMultiome data from WT and *Runx1* KO CFs at 7D and 4W post-MI demonstrated co-accessibility between Runx1-bound enhancers and promoters of target genes, such as *Comp* and *Cemip* (**Figures 6J, S6H-O**). Reduced accessibility at these enhancers was observed in *Runx1* KO MaFs and CMFs at 7D post-MI (**Figures 6J, S6N**).

Finally, gain-of-function experiments showed that *Runx1* overexpression increased *Comp* expression (**Figure 6K**). Surprisingly, *Runx1* overexpression did not elevate other tested targets (*Cdh2*, *Cemip*, and *Lox*), possibly due to the lack of co-overexpression of Runx1 co-factors required for their activation (**Figure S6P**). Together, these findings indicate that Runx1 regulates CF gene expression through direct enhancer binding and chromatin remodeling.

### *Runx1* regulates CF-macrophage communication after MI

The potential role of *Runx1* in immune and defense responses within CFs suggested that fibroblast-specific *Runx1* KO might indirectly affect immune cells (**Figures S3F, S6B**). To explore this, we reclustered Mo/Ma from 7D post-MI hearts of WT and *Runx1* KO mice. Nine clusters were identified, including monocytes (C4 Mo, *Vcan*^+^), tissue-resident macrophages (C3 Ma-3, *Cd163^+^*), and seven additional macrophage clusters likely derived from monocytes (**Figures 7A–C**). Notably, C5 Ma-4 was reduced in *Runx1* KO hearts compared to WT (**Figures 7A–B**). C5 Ma-4 expressed high levels of pro-inflammatory chemokines and cytokines (*Cxcl1*, *Cxcl10*, *Ccl4*, *Tnf*) but also the anti-inflammatory cytokine, *Il10*, resembling M2b macrophages (**Figures 7C–D**)^31^. GSEA confirmed enrichment of cytokine activity and inflammatory response terms in this cluster, indicating a strong pro-inflammatory identity (**Figure 7E**). In contrast, C1 Ma-1, which was more abundant in *Runx1* KO hearts, showed negative enrichment for these terms, suggesting a less inflammatory phenotype (**Figure 7E**).

**Figure 7.**
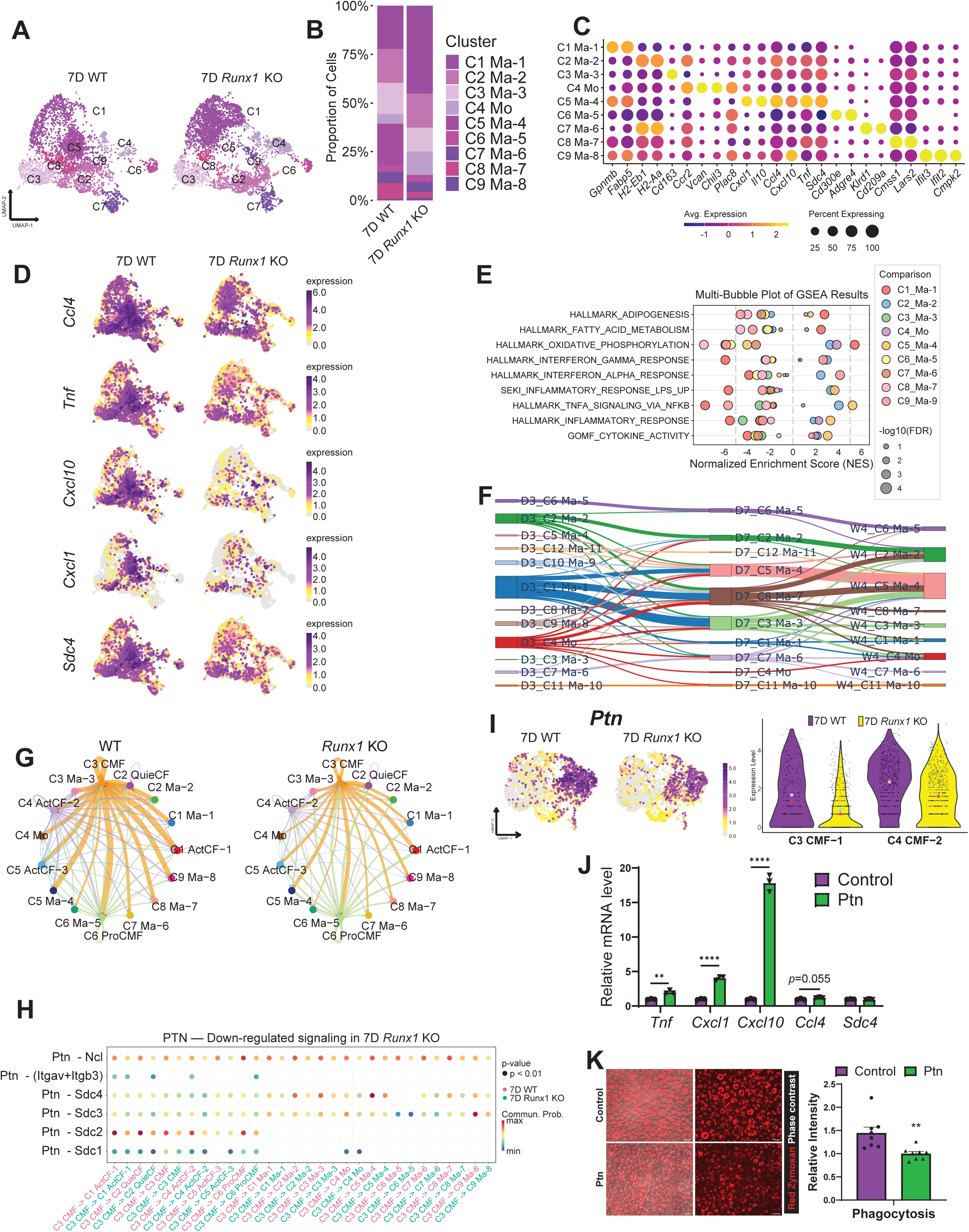
*Runx1* regulates CF-macrophage communication after MI. (A-B) UMAP graphs (A) and a bar graph (B) show different Mo/Ma subtypes identified from WT and *Runx1* KO hearts at 7D post-MI. (C-D) A dotplot (C) and feature plots (D) show the expression of selected Mo/Ma subtype marker genes at 7D post-MI. (E) A multi-bubble plot illustrates differential enrichment of GOBP terms among Mo/Ma subtypes at 7D post-MI. (F) A Sankey graph depicts the differentiation trajectory of Mo/Ma in WT hearts across indicated post-MI time points. (G) Circle plots show Ptn-mediated signaling strength between CF and Mo/Ma clusters in WT and *Runx1* KO hearts at 7D post-MI. (H) A dot plot shows Ptn-mediated communication from CMFs to Mo/Ma and other CF subtypes in WT and *Runx1* KO hearts at 7D post-MI. (I) Feature and violin plots display *Ptn* expression in WT and *Runx1* KO CFs at 7D post-MI. Triangles and dots in the violin plot indicate median and mean, respectively. (J) A bar graph shows the expression of selected genes in BMDMs treated or untreated with recombinant Ptn. (K) Images and quantification show phagocytic activities of control and Ptn-treated BMDMs. Scale bar: 35 μm. *, *p* < 0.05. n=4.

Reclustering Mo/Ma across 3D, 7D, and 4W post-MI revealed that C3 Ma-3 and C8 Ma-7 corresponded to C1 Ma-1 and C5 Ma-4 from the 7D analysis (**Figures S7A–D**). Although C8 Ma-7 declined sharply by 4W, it remained more abundant in WT hearts than in *Runx1* KO hearts (**Figure S7C**). Trajectory analysis using MOSCOT indicated that in WT hearts, C8 Ma-7 at 7D likely originated from C1 Ma-1, C4 Mo, and C10 Ma-9 at 3D and subsequently gave rise to C2 Ma-2 and C5 Ma-4 at 4W (**Figure 7F**). Pseudotime analysis using CellRank2 supported a monocyte origin for C8 Ma-7, as both C1 Ma-1 and C10 Ma-9 traced back to C4 Mo (**Figure S7E**). In *Runx1* KO hearts, C8 Ma-7 appeared to be largely replaced by C3 Ma-3 (**Figure S7F**). GSEA showed that C8 Ma-7 was strongly enriched for pro-inflammatory pathways compared to its ancestor clusters (C1 Ma-1, C4 Mo, C10 Ma-9), descendant clusters (C2 Ma-2, C5 Ma-4), and parallel cluster (C3 Ma-3) (**Figure S7G**). These findings suggest that WT hearts exhibit a transient surge in pro-inflammatory macrophages at 7D post-MI, which is diminished in *Runx1* KO hearts.

To investigate the mechanisms underlying these differences, we performed cell-cell communication analysis on 7D post-MI scRNAseq data using CellChat^32^. The analysis revealed robust interactions between CF clusters and macrophage clusters, with Ptn-Sdc4 signaling between CMF (CMF-1 + CMF-2) and Ma-4 showing the greatest upregulation in WT compared to *Runx1* KO hearts (**Figures 7G-H, S7H**). *Ptn* was identified as a Runx1 target gene in the GRN and was expressed at higher levels in WT versus *Runx1* KO C4 CMF-2 (**Figure 7I; Table S9**). *Runx1* overexpression further increased *Ptn* expression in CFs (**Figure S7I**). To test whether Ptn mediates macrophage activity, bone marrow-derived macrophages (BMDMs) were treated with recombinant Ptn, which significantly increased expression of chemokines and cytokines enriched in C5 Ma-4 but not the receptor gene *Sdc4* (**Figure 7J**). Interestingly, Ptn treatment reduced the phagocytic activity of BMDMs (**Figure 7K**). These results indicate that *Runx1* in CFs regulates macrophage inflammatory cytokine secretion through *Ptn*-mediated signaling, which inhibits the ability of macrophages to clear tissue debris and likely hampers tissue repair. Interestingly, although *Ptn* expression and associated epigenetic marks were elevated in CFs at 7D post-MI and in cultured WT CMFs compared to *Runx1* KO CMFs, Runx1 binding in regulatory elements within or near the *Ptn* locus was less evident, suggesting unstable interactions (**Figures S7J-K**).

### Combined *Runx1*/*Runx2* KO amplifies the effects of *Runx1* Loss

The Runx transcription factor family includes *Runx1*, *Runx2*, and *Runx3*. In addition to *Runx1*, *Runx2* is active in CFs (**Figures S8A-B**). Approximately half of Runx2 targets identified by SCENIC+ overlapped with Runx1 targets, and Runx2+ regulon enrichment was observed in WT versus *Runx1* KO CMFs at 7D post-MI (**Figures S8C-D**). The pronounced impact of *Runx1* KO on C4 ActCF-2 at 4W post-MI is likely attributed to the absence of *Runx2* expression and its compensatory effect in this cluster (**Figures 4M-P**). These results suggest partial functional redundancy between *Runx1* and *Runx2*.

To test whether *Runx2* deletion augments *Runx1* KO effects, we generated fibroblast-specific *Runx1*/*Runx2* double KO mice (*Runx1/2* KO) and performed scRNAseq at 7D post-MI (**Figures S8E-F**). Reclustering revealed a more severe disruption of CF subtype composition in *Runx1/2* KO hearts compared to *Runx1* KO, with a further increase in QuieCFs (C2) and a greater reduction in the CMF-2/CMF-1 ratio (**Figures 8A-C**). Runx1-targted fibrotic and chondrogenic genes, including *Postn*, *Acta2*, *Ptn*, *Comp*, *Cthrc1*, *Acan*, *Col1a1*, *Col11a1*, and *Col2a1*, and relevant GOBP term enrichment showed stronger reductions in *Runx1/2* KO than in *Runx1* KO CMFs versus WT (**Figures 8D-F, Tables S15-S16**). Conversely, *Pdgfra*, a QuieCF marker, was upregulated in *Runx1/2* KO CMFs compared to WT and *Runx1* KO, indicating that combined deletion strongly inhibits CF activation and differentiation (**Figures 8D-E**). Consistently, enrichment of Runx1+ and Runx2+ regulons was further diminished in double KO vs *Runx1* KO CMFs (**Figure S8D**). Pseudotime analysis confirmed that the differentiation trajectory from CMF-1 to CMF-2 was markedly impaired in *Runx1/2* KO hearts (**Figures 8C, S8G**).

**Figure 8.**
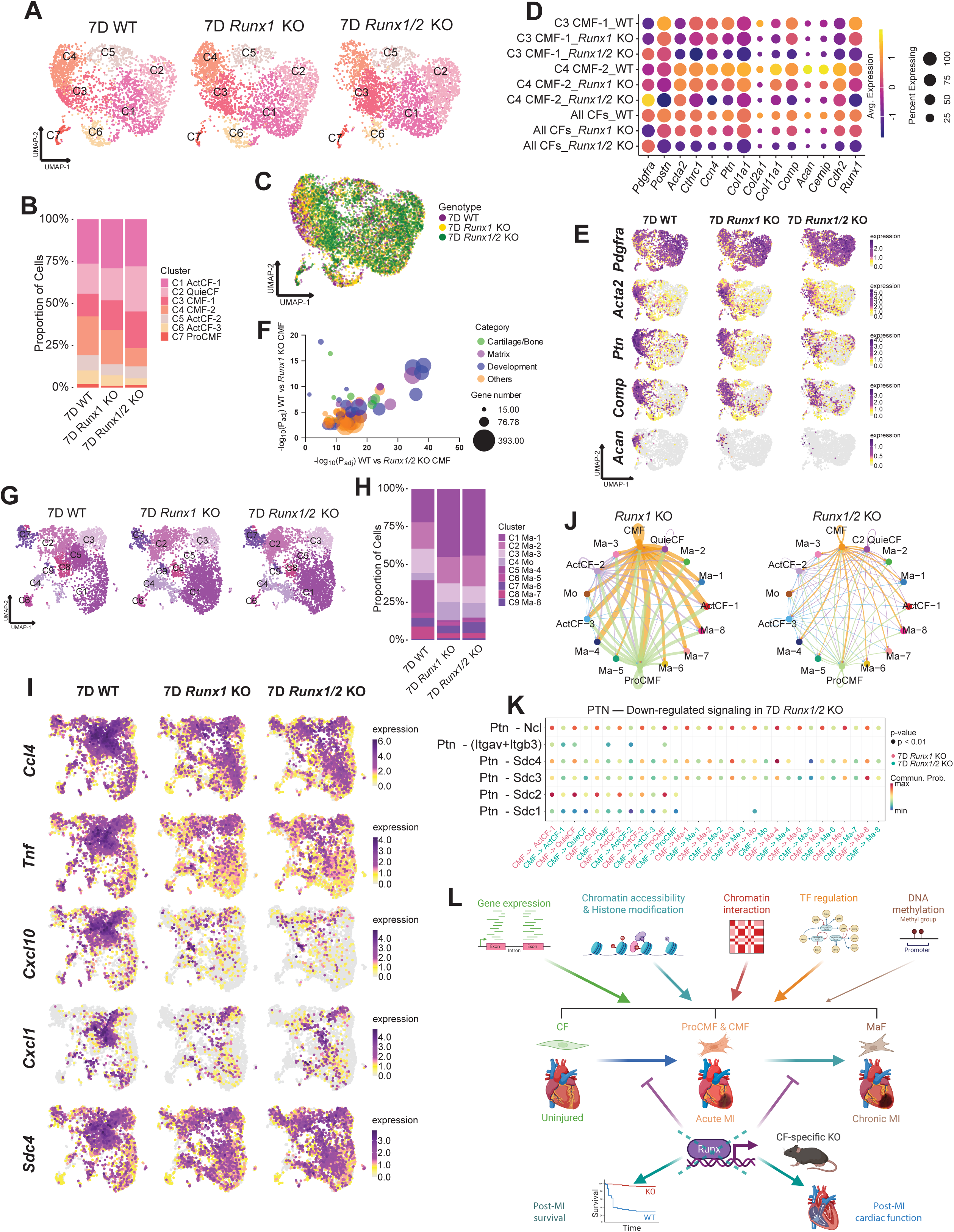
Combined *Runx1* and *Runx2* KO enhances the effects of *Runx1* deletion in CFs. (A-B) UMAP graphs (A) and a bar graph (B) show CF subtypes identified in WT, *Runx1* KO, and *Runx1/2* KO hearts at 7D post-MI. (C) A UMAP graph shows CFs identified in WT, *Runx1* KO, and *Runx1/2* KO hearts at 7D post-MI, with cells color-coded by genotypes. (D) A dot plot shows expression of selected genes in C3 CMF-1, C4 CMF-2, and all CFs across WT, *Runx1* KO, and *Runx1/2* KO hearts at 7D post-MI. (E) Feature plots display expression of selected genes in CFs from WT, *Runx1* KO, and *Runx1/2* KO hearts at 7D post-MI. (F) A multivariable plot shows differential enrichment of GOBP terms related to cartilage/bone formation, extracellular matrix, and development between WT and *Runx1* KO CMFs and between WT and *Runx1/2* KO CMFs. (G-H) UMAP graphs (G) and a bar graph (H) show Mo/Ma subtypes identified in WT, *Runx1* KO, and *Runx1/2* KO hearts at 7D post-MI. (I) Feature plots display the expression of selected genes in Mo/Ma subtypes across WT, *Runx1* KO, and *Runx1/2* KO hearts at 7D post-MI. (J) Circle plots show Ptn-mediated signaling between CF and Mo/Ma clusters in *Runx1* KO and *Runx1/2* KO hearts at 7D post-MI. (K) A dot plot shows Ptn-mediated communication from CMFs to CF and Mo/Ma subtypes in *Runx1* KO and *Runx1/2* KO hearts at 7D post-MI. (L) A graphic summary of this study.

To assess indirect effects on immune cells, we reclustered Mo/Ma populations from WT, *Runx1* KO, and *Runx1/2* KO hearts at 7D post-MI (**Figures 8G-H**). The Mo/Ma composition in *Runx1/2* KO hearts largely mirrored that of *Runx1* KO hearts, with reduced abundance of pro-inflammatory C5 Ma-4 and increased abundance of C1 Ma-1 (**Figures 8G-I**). Cell-cell communication analysis revealed further diminished Ptn–Sdc4 signaling between CMFs and Ma-4 in *Runx1/2* KO hearts (**Figures 8J-K, S8H-I**).

These findings demonstrate that combined *Runx1*/*Runx2* deletion amplifies the effects of *Runx1* loss, leading to stronger suppression of CF activation and differentiation and further weakening CF–macrophage interactions.

## DISCUSSION

Although CFs are known to play a central role in post-MI fibrosis, a comprehensive analysis of their dynamics throughout the entire remodeling process has been lacking^33–35^. Here, we provide a high-resolution view of CF gene expression using integrated single-nucleus and bulk transcriptomic approaches. CFs were rapidly activated within hours of MI, as early as 2H post-injury, marked by increased expression of genes involved in transcription, translation, and proliferation, well before cell division begins. This early activation likely primes CFs for the robust proliferation observed around day 3, which is essential for maintaining infarct structural integrity. Unlike macrophages and vascular cells, which retain proliferative populations up to 4W post-MI, CF proliferation declines soon after day 3, potentially limiting excessive fibrosis. By day 7, CFs in the infarct expressed both CMF and MaF signatures, indicating ongoing differentiation and revealing that the CMF state is more transient than previously thought^10^, making clear boundaries between CF states difficult to define.

A second major focus of this study was multiomic profiling of CFs. Despite the recognized importance of epigenetic regulation, systematic analysis of its role in CF activity after MI has been limited by technical challenges. We addressed this by combining single-cell/nucleus and bulk sequencing with lineage tracing to profile gene expression, chromatin accessibility, histone modifications, chromatin interactions, and DNA methylation. Gene expression changes closely correlated with alterations in chromatin accessibility and histone marks, often involving promoter– enhancer loops (**Figure 8L**). In contrast, DNA methylation contributed minimally to MI-induced transcriptomic changes, instead correlating with transcriptional activity variations among genes at individual time points (**Figure 8L**). This supports the view that DNA methylation provides stable lineage-specific silencing, while the more reversible histone modifications play a major role in transient gene expression regulation^36^. Notably, although H3K27me3 levels negatively correlated with gene expression, they showed no association with DNA methylation, suggesting that polycomb repressive complex 2 and DNA methylation act independently in CFs, despite reported crosstalk in embryonic stem cell differentiation^37, 38^.

We further refined the SCENIC+ pipeline to incorporate TADs and histone modification– inferred enhancer activity for regulon identification. Standard SCENIC+ does not account for long-range enhancer interactions or enhancer activity changes not reflected in chromatin accessibility. Our modifications enabled identification of additional TF targets and construction of more complete GRNs. Using this approach, we identified Runx1 as a key regulator of CF activation and differentiation after MI. Although *Runx1* has been implicated in CF biology, its precise role in post-MI remodeling was unclear^16, 39^. Analysis of Runx1 targets revealed a dual function, promoting CF proliferation and MaF differentiation. Here, we show that *Runx1* facilitates chromatin remodeling and regulates gene expression through enhancer targeting. Moreover, *in vivo*, fibroblast-specific *Runx1* KO reduced CF proliferation at 3D and impaired CMF maturation and MaF differentiation at 7D, but had little effect on fully differentiated MaFs at 4W. These findings suggest that *Runx1* accelerates the QuieCF-CMF-MaF transition, consistent with the transient nature of CMFs (**Figure 8L**).

Beyond CF-autonomous effects, *Runx1* in CFs also influenced macrophage activity by inducing inflammatory cytokines, potentially via Ptn and macrophage-expressed Sdc4. Enhanced debris clearing efficiency of macrophages, combined with reduced CF activation, may contribute to the lower incidence of cardiac rupture and improved remodeling observed in *Runx1* KO mice after MI (**Figure 8L**).

Finally, we observed functional redundancy between *Runx1* and *Runx2*, which share many targets. Combined *Runx1/2* KO caused more severe disruption of CF differentiation and CF–macrophage communication than *Runx1* KO alone. Future studies are needed to clarify their distinct and overlapping roles and assess whether double KO offers greater therapeutic benefit.

In summary, our study reveals dynamic CF activities after MI, governed by chromatin remodeling and key TFs such as Runx family members. Fibroblast-specific *Runx1* KO improved survival and reduced maladaptive remodeling, highlighting Runx TFs as potential therapeutic targets. The multiomic CF dataset generated here provides a valuable resource for further investigations.

## AUTHOR CONTRIBUTIONS

X.F. conceived the study and supervised the entire work. J.S. co-supervised sequencing data analyses. Y.L., L.W., X.Z., K.G., S.U., and B.L. performed experiments. Y.L., L.W., X.Z., K.G., N.K., M.S., R.S., R.L., J.S. and X.F. analyzed data. Y.L., J.F., C.A.S, J.S, and X.F interpreted the data, assembled the results, and wrote the manuscript with input from all authors.

## ACKNOWLEDGMENTS

The authors thank Louisiana State University (LSU) Gene Lab for help with snMultiome and scRNAseq library construction, LSU Flow Cytometry Core for help with FACS, LSU Shared Instrument Facility for help with confocal imaging, and LSU and ODU High Performance Computing for assistance with sequencing data analysis. MIGR1 was a gift from Warren Pear (Addgene plasmid # 27490; http://n2t.net/addgene:27490; RRID:Addgene_27490).

## Supplemental Material

## REFERENCES

1. Tsao CW, Aday AW, Almarzooq ZI, Alonso A, Beaton AZ, Bittencourt MS, Boehme AK, Buxton AE, Carson AP, Commodore-Mensah Y, Elkind MSV, Evenson KR, Eze-Nliam C, Ferguson JF, Generoso G, Ho JE, Kalani R, Khan SS, Kissela BM, Knutson KL, Levine DA, Lewis TT, Liu J, Loop MS, Ma J, Mussolino ME, Navaneethan SD, Perak AM, Poudel R, Rezk-Hanna M, Roth GA, Schroeder EB, Shah SH, Thacker EL, VanWagner LB, Virani SS, Voecks JH, Wang NY, Yaffe K, Martin SS. Heart Disease and Stroke Statistics-2022 Update: A Report From the American Heart Association. Circulation. 2022;145(8):e153–e639. Epub 20220126. doi: 10.1161/cir.0000000000001052. PubMed PMID: 35078371.

2. Priebe H-J. Perioperative myocardial infarction—aetiology and prevention. British journal of anaesthesia. 2005;95(1):3–19. PubMed PMID: 15665072.

3. Oka T, Xu J, Kaiser RA, Melendez J, Hambleton M, Sargent MA, Lorts A, Brunskill EW, Dorn GW, 2nd, Conway SJ, Aronow BJ, Robbins J, Molkentin JD. Genetic manipulation of periostin expression reveals a role in cardiac hypertrophy and ventricular remodeling. Circ Res. 2007;101(3):313–21. Epub 20070614. doi: 10.1161/circresaha.107.149047. PubMed PMID: 17569887; PMCID: PMC2680305.

4. Trindade ML, Tsutsui JM, Rodrigues AC, Caldas MA, Ramires JA, Mathias Junior W. Left ventricular free wall impeding rupture in post-myocardial infarction period diagnosed by myocardial contrast echocardiography: case report. Cardiovasc Ultrasound. 2006;4:7. Epub 20060126. doi: 10.1186/1476-7120-4-7. PubMed PMID: 16438720; PMCID: PMC1395330.

5. Kong P, Christia P, Frangogiannis NG. The pathogenesis of cardiac fibrosis. Cell Mol Life Sci. 2014;71(4):549–74. Epub 20130507. doi: 10.1007/s00018-013-1349-6. PubMed PMID: 23649149; PMCID: PMC3769482.

6. Janicki JS, Brower GL. The role of myocardial fibrillar collagen in ventricular remodeling and function. J Card Fail. 2002;8(6 Suppl):S319-25. doi: 10.1054/jcaf.2002.129260. PubMed PMID: 12555139.

7. Kanisicak O, Khalil H, Ivey MJ, Karch J, Maliken BD, Correll RN, Brody MJ, SC JL, Aronow BJ, Tallquist MD, Molkentin JD. Genetic lineage tracing defines myofibroblast origin and function in the injured heart. Nat Commun. 2016;7:12260. Epub 20160722. doi: 10.1038/ncomms12260. PubMed PMID: 27447449; PMCID: PMC5512625.

8. Humeres C, Frangogiannis NG. Fibroblasts in the Infarcted, Remodeling, and Failing Heart. JACC Basic Transl Sci. 2019;4(3):449–67. Epub 20190624. doi: 10.1016/j.jacbts.2019.02.006. PubMed PMID: 31312768; PMCID: PMC6610002.

9. Ivey MJ, Kuwabara JT, Pai JT, Moore RE, Sun Z, Tallquist MD. Resident fibroblast expansion during cardiac growth and remodeling. J Mol Cell Cardiol. 2018;114:161–74. Epub 20171120. doi: 10.1016/j.yjmcc.2017.11.012. PubMed PMID: 29158033; PMCID: PMC5831691.

10. Fu X, Khalil H, Kanisicak O, Boyer JG, Vagnozzi RJ, Maliken BD, Sargent MA, Prasad V, Valiente-Alandi I, Blaxall BC, Molkentin JD. Specialized fibroblast differentiated states underlie scar formation in the infarcted mouse heart. J Clin Invest. 2018;128(5):2127–43. Epub 20180416. doi: 10.1172/jci98215. PubMed PMID: 29664017; PMCID: PMC5957472.

11. Fortier SM, Penke LR, King D, Pham TX, Ligresti G, Peters-Golden M. Myofibroblast dedifferentiation proceeds via distinct transcriptomic and phenotypic transitions. JCI Insight. 2021;6(6). Epub 20210322. doi: 10.1172/jci.insight.144799. PubMed PMID: 33561015; PMCID: PMC8026183.

12. Desmoulière A, Redard M, Darby I, Gabbiani G. Apoptosis mediates the decrease in cellularity during the transition between granulation tissue and scar. Am J Pathol. 1995;146(1):56–66. PubMed PMID: 7856739; PMCID: PMC1870783.

13. Amrute JM, Luo X, Penna V, Yang S, Yamawaki T, Hayat S, Bredemeyer A, Jung IH, Kadyrov FF, Heo GS, Venkatesan R, Shi SY, Parvathaneni A, Koenig AL, Kuppe C, Baker C, Luehmann H, Jones C, Kopecky B, Zeng X, Bleckwehl T, Ma P, Lee P, Terada Y, Fu A, Furtado M, Kreisel D, Kovacs A, Stitziel NO, Jackson S, Li CM, Liu Y, Rosenthal NA, Kramann R, Ason B, Lavine KJ. Targeting immune-fibroblast cell communication in heart failure. Nature. 2024;635(8038):423–33. Epub 20241023. doi: 10.1038/s41586-024-08008-5. PubMed PMID: 39443792; PMCID: PMC12334188.

14. Alexanian M, Padmanabhan A, Nishino T, Travers JG, Ye L, Pelonero A, Lee CY, Sadagopan N, Huang Y, Auclair K, Zhu A, An Y, Ekstrand CA, Martinez C, Teran BG, Flanigan WR, Kim CK, Lumbao-Conradson K, Gardner Z, Li L, Costa MW, Jain R, Charo I, Combes AJ, Haldar SM, Pollard KS, Vagnozzi RJ, McKinsey TA, Przytycki PF, Srivastava D. Chromatin remodelling drives immune cell-fibroblast communication in heart failure. Nature. 2024;635(8038):434–43. Epub 20241023. doi: 10.1038/s41586-024-08085-6. PubMed PMID: 39443808; PMCID: PMC11698514.

15. Vasquez C, Mohandas P, Louie KL, Benamer N, Bapat AC, Morley GE. Enhanced fibroblast-myocyte interactions in response to cardiac injury. Circ Res. 2010;107(8):1011–20. Epub 20100812. doi: 10.1161/circresaha.110.227421. PubMed PMID: 20705922; PMCID: PMC2993566.

16. Li C, Sun J, Liu Q, Dodlapati S, Ming H, Wang L, Li Y, Li R, Jiang Z, Francis J, Fu X. The landscape of accessible chromatin in quiescent cardiac fibroblasts and cardiac fibroblasts activated after myocardial infarction. Epigenetics. 2022;17(9):1020–39. Epub 20211025. doi: 10.1080/15592294.2021.1982158. PubMed PMID: 34551670; PMCID: PMC9487753.

17. Alexanian M, Przytycki PF, Micheletti R, Padmanabhan A, Ye L, Travers JG, Gonzalez-Teran B, Silva AC, Duan Q, Ranade SS, Felix F, Linares-Saldana R, Li L, Lee CY, Sadagopan N, Pelonero A, Huang Y, Andreoletti G, Jain R, McKinsey TA, Rosenfeld MG, Gifford CA, Pollard KS, Haldar SM, Srivastava D. A transcriptional switch governs fibroblast activation in heart disease. Nature. 2021;595(7867):438–43. Epub 20210623. doi: 10.1038/s41586-021-03674-1. PubMed PMID: 34163071; PMCID: PMC8341289.

18. Stratton MS, Bagchi RA, Felisbino MB, Hirsch RA, Smith HE, Riching AS, Enyart BY, Koch KA, Cavasin MA, Alexanian M, Song K, Qi J, Lemieux ME, Srivastava D, Lam MPY, Haldar SM, Lin CY, McKinsey TA. Dynamic Chromatin Targeting of BRD4 Stimulates Cardiac Fibroblast Activation. Circ Res. 2019;125(7):662–77. Epub 20190814. doi: 10.1161/circresaha.119.315125. PubMed PMID: 31409188; PMCID: PMC7310347.

19. Small EM, Thatcher JE, Sutherland LB, Kinoshita H, Gerard RD, Richardson JA, Dimaio JM, Sadek H, Kuwahara K, Olson EN. Myocardin-related transcription factor-a controls myofibroblast activation and fibrosis in response to myocardial infarction. Circ Res. 2010;107(2):294–304. Epub 20100617. doi: 10.1161/circresaha.110.223172. PubMed PMID: 20558820; PMCID: PMC2921870.

20. Xiao Y, Hill MC, Li L, Deshmukh V, Martin TJ, Wang J, Martin JF. Hippo pathway deletion in adult resting cardiac fibroblasts initiates a cell state transition with spontaneous and self-sustaining fibrosis. Genes Dev. 2019;33(21-22):1491–505. Epub 20190926. doi: 10.1101/gad.329763.119. PubMed PMID: 31558567; PMCID: PMC6824468.

21. Olsen MB, Hildrestrand GA, Scheffler K, Vinge LE, Alfsnes K, Palibrk V, Wang J, Neurauter CG, Luna L, Johansen J, Øgaard JDS, Ohm IK, Slupphaug G, Kuśnierczyk A, Fiane AE, Brorson SH, Zhang L, Gullestad L, Louch WE, Iversen PO, Østlie I, Klungland A, Christensen G, Sjaastad I, Sætrom P, Yndestad A, Aukrust P, Bjørås M, Finsen AV. NEIL3-Dependent Regulation of Cardiac Fibroblast Proliferation Prevents Myocardial Rupture. Cell Rep. 2017;18(1):82–92. doi: 10.1016/j.celrep.2016.12.009. PubMed PMID: 28052262.

22. Chung MI, Bujnis M, Barkauskas CE, Kobayashi Y, Hogan BLM. Niche-mediated BMP/SMAD signaling regulates lung alveolar stem cell proliferation and differentiation. Development. 2018;145(9). Epub 20180511. doi: 10.1242/dev.163014. PubMed PMID: 29752282; PMCID: PMC5992594.

23. Yamamoto M, Shook NA, Kanisicak O, Yamamoto S, Wosczyna MN, Camp JR, Goldhamer DJ. A multifunctional reporter mouse line for Cre- and FLP-dependent lineage analysis. Genesis. 2009;47(2):107–14. Epub 2009/01/24. doi: 10.1002/dvg.20474. PubMed PMID: 19165827; PMCID: PMC8207679.

24. Takarada T, Hinoi E, Nakazato R, Ochi H, Xu C, Tsuchikane A, Takeda S, Karsenty G, Abe T, Kiyonari H, Yoneda Y. An analysis of skeletal development in osteoblast-specific and chondrocyte-specific runt-related transcription factor-2 (Runx2) knockout mice. J Bone Miner Res. 2013;28(10):2064–9. doi: 10.1002/jbmr.1945. PubMed PMID: 23553905.

25. Thompson SA, Blazeski A, Copeland CR, Cohen DM, Chen CS, Reich DM, Tung L. Acute slowing of cardiac conduction in response to myofibroblast coupling to cardiomyocytes through N-cadherin. J Mol Cell Cardiol. 2014;68:29–37. Epub 20140109. doi: 10.1016/j.yjmcc.2013.12.025. PubMed PMID: 24412534; PMCID: PMC3993970.

26. Bravo González-Blas C, De Winter S, Hulselmans G, Hecker N, Matetovici I, Christiaens V, Poovathingal S, Wouters J, Aibar S, Aerts S. SCENIC+: single-cell multiomic inference of enhancers and gene regulatory networks. Nat Methods. 2023;20(9):1355–67. Epub 20230713. doi: 10.1038/s41592-023-01938-4. PubMed PMID: 37443338; PMCID: PMC10482700.

27. Hafemeister C, Satija R. Normalization and variance stabilization of single-cell RNA-seq data using regularized negative binomial regression. Genome Biol. 2019;20(1):296. Epub 20191223. doi: 10.1186/s13059-019-1874-1. PubMed PMID: 31870423; PMCID: PMC6927181.

28. Weiler P, Lange M, Klein M, Pe’er D, Theis F. CellRank 2: unified fate mapping in multiview single-cell data. Nat Methods. 2024;21(7):1196–205. Epub 20240613. doi: 10.1038/s41592-024-02303-9. PubMed PMID: 38871986; PMCID: PMC11239496.

29. Sekiya I, Vuoristo JT, Larson BL, Prockop DJ. In vitro cartilage formation by human adult stem cells from bone marrow stroma defines the sequence of cellular and molecular events during chondrogenesis. Proc Natl Acad Sci U S A. 2002;99(7):4397–402. Epub 20020326. doi: 10.1073/pnas.052716199. PubMed PMID: 11917104; PMCID: PMC123659.

30. Acharya A, Baek ST, Banfi S, Eskiocak B, Tallquist MD. Efficient inducible Cre-mediated recombination in Tcf21 cell lineages in the heart and kidney. Genesis. 2011;49(11):870–7. Epub 2011/03/25. doi: 10.1002/dvg.20750. PubMed PMID: 21432986; PMCID: PMC3279154.

31. Wang LX, Zhang SX, Wu HJ, Rong XL, Guo J. M2b macrophage polarization and its roles in diseases. J Leukoc Biol. 2019;106(2):345–58. Epub 20181221. doi: 10.1002/jlb.3ru1018-378rr. PubMed PMID: 30576000; PMCID: PMC7379745.

32. Jin S, Plikus MV, Nie Q. CellChat for systematic analysis of cell-cell communication from single-cell transcriptomics. Nat Protoc. 2025;20(1):180–219. Epub 20240916. doi: 10.1038/s41596-024-01045-4. PubMed PMID: 39289562.

33. Tallquist MD, Molkentin JD. Redefining the identity of cardiac fibroblasts. Nat Rev Cardiol. 2017;14(8):484–91. Epub 20170424. doi: 10.1038/nrcardio.2017.57. PubMed PMID: 28436487; PMCID: PMC6329009.

34. Venugopal H, Hanna A, Humeres C, Frangogiannis NG. Properties and Functions of Fibroblasts and Myofibroblasts in Myocardial Infarction. Cells. 2022;11(9). Epub 20220420. doi: 10.3390/cells11091386. PubMed PMID: 35563692; PMCID: PMC9102016.

35. Daseke MJ, 2nd, Tenkorang MAA, Chalise U, Konfrst SR, Lindsey ML. Cardiac fibroblast activation during myocardial infarction wound healing: Fibroblast polarization after MI. Matrix Biol. 2020;91–92:109-16. Epub 20200521. doi: 10.1016/j.matbio.2020.03.010. PubMed PMID: 32446909; PMCID: PMC7434699.

36. Cedar H, Bergman Y. Linking DNA methylation and histone modification: patterns and paradigms. Nat Rev Genet. 2009;10(5):295–304. doi: 10.1038/nrg2540. PubMed PMID: 19308066.

37. McLaughlin K, Flyamer IM, Thomson JP, Mjoseng HK, Shukla R, Williamson I, Grimes GR, Illingworth RS, Adams IR, Pennings S, Meehan RR, Bickmore WA. DNA Methylation Directs Polycomb-Dependent 3D Genome Re-organization in Naive Pluripotency. Cell Rep. 2019;29(7):1974–85.e6. doi: 10.1016/j.celrep.2019.10.031. PubMed PMID: 31722211; PMCID: PMC6856714.

38. Mohn F, Weber M, Rebhan M, Roloff TC, Richter J, Stadler MB, Bibel M, Schübeler D. Lineage-specific polycomb targets and de novo DNA methylation define restriction and potential of neuronal progenitors. Mol Cell. 2008;30(6):755–66. Epub 20080529. doi: 10.1016/j.molcel.2008.05.007. PubMed PMID: 18514006.

39. Amrute JM, Lai L, Ma P, Koenig AL, Kamimoto K, Bredemeyer A, Shankar TS, Kuppe C, Kadyrov FF, Schulte LJ, Stoutenburg D, Kopecky BJ, Navankasattusas S, Visker J, Morris SA, Kramann R, Leuschner F, Mann DL, Drakos SG, Lavine KJ. Defining cardiac functional recovery in end-stage heart failure at single-cell resolution. Nat Cardiovasc Res. 2023;2(4):399–416. Epub 20230406. doi: 10.1038/s44161-023-00260-8. PubMed PMID: 37583573; PMCID: PMC10426763.

