## Supplemental Methods and Figures for "Multiomics reveals epigenetic control of fibroblast activity after myocardial infarction and a key role for RUNX transcription factors"

**Supplemental Material**

Supplemental Methods

Tables S1–S15

Figure S1-S8

### Supplemental Methods

#### Animals

$Pdgfra^{CreERT2}$ mice (JAX:032770)[1] were crossed with $R26^{GFP}$ mice[2] to generate $Pdgfra^{CreERT2}$;$R26^{GFP}$ mice. To generate $Pdgfra^{CreERT2}$;$Runx1^{fl/fl}$;$R26^{GFP}$ and $Pdgfra^{CreERT2}$;$Runx1^{fl/fl}$;$Runx2^{fl/fl}$;$R26^{GFP}$ mice, $Pdgfra^{CreERT2}$;$R26^{GFP}$ mice were crossed with $Runx1^{fl/fl}$ (JAX:010673) or/and $Runx2^{fl/fl}$[3] mice. All mice are in C57BL/6J background. All experiments involving mice were approved by the IACUC at LSU.

#### Tamoxifen treatment in mice

To induce the activity of Cre, mice were given tamoxifen (Sigma, #T5648) dissolved in corn oil *via* intraperitoneal (i.p.) injections or gavage at approximately 6 weeks of age at a dosage of 75 mg/kg body weight /day for 5 consecutive days.

#### Myocardial infarction

Mice were subjected to permanent surgical ligation of the left coronary artery to induce MI at 9 weeks of age using our established protocol. Briefly, mice were anesthetized using isoflurane by a ventilator at 1-3% throughout the entire course of the procedure. A left lateral thoracotomy was performed. The left anterior descending coronary artery was identified and ligated just below the left atrium. For pain management related to surgical procedures, mice were given a dose of Carprofen (5 mg/kg body weight) by subcutaneous injection before the surgery followed by a second dose 12 hours after the surgery. Mice were checked every day after MI for survival rate. Mice found dead were subjected to necropsy to identify cardiac rupture. Mice were euthanized by isoflurane overdose followed by cervical dislocation before sample collections.

#### *In vivo* EdU fibroblast proliferation assay

At 3D and 7D post-MI, mice were treated with EdU through i.p. injections at a dose of 50 mg/kg body weight. Four hours after EdU administration, the mice were euthanized, and heart tissues were collected. EdU incorporation was detected by immunohistochemical staining using the Click-&-Go Plus EdU 647 Cell Proliferation Assay Kit (Click Chemistry Tools, #1353).

#### Echocardiography

Echocardiography was conducted before left anterior descending coronary artery ligation for baseline cardiac function and at 7D and 4W post-MI using a Vevo F2 LT Imaging System equipped with UHF46x transducer (FUJIFILM Visual Sonics, Inc.). Mice were initially anesthetized with 5% isoflurane and 1L/minute medical oxygen, then maintained under light anesthesia with 1–2% isoflurane via a nose cone on a heated platform. Heart rate was continuously monitored and maintained between 400-500 bpm by securing the mice's paws to the electrodes on the platform using dermatological tape during imaging. After dehairing, echocardiographic imaging was performed following standard protocols, acquiring parasternal long-axis and short-axis views in both B-mode and M-mode. Cardiac function, including ejection fraction (EF) and fractional shortening (FS), was evaluated using Simpson's method based on left ventricular long-axis B-mode imaging and three short-axis B-mode images (proximal, mid, and distal).

#### Tissue processing for histological analysis

Hearts were fixed in 4% paraformaldehyde (PFA) (Electron Microscopy Sciences, #15714) in PBS at 4°C for 4 hours, followed by three 5-minute washes in PBS. Samples were then incubated in PBS containing 30% sucrose at 4°C overnight and subsequently equilibrated in a 1:1 mixture of OCT and 30% sucrose for 30 minutes. Following equilibration, tissues were embedded in OCT (SAKURA, #4583) within cryomolds and snap-frozen in the liquid nitrogen-chilled isopentane for 20 seconds with gentle agitation. Frozen tissues were immediately transferred to a −80°C freezer until sectioning. Cryosections at 5μm thickness were prepared at the desired cryotome temperature and air-dried at room temperature for 30 minutes. Slides were either processed immediately for staining or stored at −80°C until further use.

**Immunocytochemical staining**

Cells were fixed in either 4% PFA in PBS or 100% ice-cold methanol for 10 minutes on ice, followed by three washes with 1× TBS for 5 minutes each. Cells were then incubated in blocking buffer (5% goat or donkey serum, 1% BSA, 0.2% Triton™ X-100, 0.1% Sodium Azide in 1x TBS) for 1 hour, after which primary antibodies, diluted in blocking buffer, were applied to the cells for overnight incubation at 4°C. Cells were washed three times with 0.2% Triton X-100 in TBS (TBST) for 5 minutes each, then incubated with fluorochrome-conjugated secondary antibodies diluted in blocking buffer for 1 hour at room temperature in the dark. Following three additional washes in TBST, cells were mounted with DAPI-containing mounting medium (Vector Laboratories, #H-1200-10). For EdU staining, a Click-&-Go Plus EdU 647 Cell Proliferation Assay Kit (Click Chemistry Tools, #1353) was used following the manufacturer's protocol after secondary antibody staining.

**Immunohistochemical staining**

Prior to staining, slides were rinsed in double-distilled water three times for 5 minutes each. An IHC PAP (mini) pen (Enzo, #ADI-950-232-0001) was used to draw a hydrophobic barrier around the cryosections. Sections were incubated in blocking buffer for 1 hour, incubated with primary antibodies diluted in blocking buffer overnight at 4°C, and then with fluorophore-conjugated secondary antibodies diluted in blocking buffer for 1 hour at room temperature. After staining, sections were mounted with DAPI-containing mounting medium (Vector Laboratories, #H-1200-10). Images were acquired using an ECHO Revolve microscope or an Olympus CSU W1 Spinning Disk confocal.

**Trichrome staining**

Masson's Trichrome staining was performed using reagents purchased from Electron Microscopy Sciences following the manufacturer's protocol. The infarct fraction was determined using Image J.

**Cardiac fibroblast isolation and culture**

Hearts were excised, rinsed three times with sterile PBS, and kept on ice. Atria and excess tissues, including blood vessels and fat, were removed using sterile scissors and forceps. The remaining tissue was minced in 300–600 μL of digestion buffer [ 0.75 U/ml collagenase D (Roche, #11088866001), 1.0 U/ml Dispase II (Roche, #10165859001), and 2 mM CaCl2 in DMEM] within a 1.5 mL microcentrifuge tube and transferred to a 5 mL tube. The volume was adjusted to 3 mL with digestion buffer (for 1–3 hearts) and incubated at 37 °C with shaking (150 rpm) for 25 minutes. Following incubation, the mixture was gently pipetted, allowed to settle for 1 minute, and the cell suspension was collected into a 50 mL tube containing 20 mL of ice-cold PBS. Residual tissue was digested again with 1.5–2 mL of buffer for 15–20 minutes under the same conditions and briefly vortexed, and the supernatant was combined with the previous fraction. The combined cell suspension was filtered sequentially through 100 μm and 40 μm strainers and centrifuged at 700 × g for 10 minutes. The pellet was resuspended in Red Blood Cell Lysis Buffer (Sigma, # R775) and mixed gently for 1–3 minutes before dilution with 15–30 mL of ice-cold PBS, followed by centrifugation at 700 × g for 10 minutes. The final cell pellet was resuspended in a complete medium (10% bovine growth serum, 50μg/ml gentamycin, 1% of an antibiotic mixture containing

10,000 U/ml penicillin, 10 mg/ml streptomycin, and 25 μg/ml amphotericin B in high glucose DMEM). If used without FACS, CFs were enriched by removing unattached cells after 1 hour of incubation. To overexpress *Runx1* in CFs, cells were treated with retrovirus expressing GFP, mouse *Runx1*, and human *RUNX1* at 20-30 multiplicity of infection (MOI) together with 8μg/ml polybrene. Myofibroblast differentiation was induced by treating cells with TGFβ (10 ng/ml) for 48 hours. For cells subjected to the EdU assay, the culture medium was replaced with medium containing 10 μM EdU 6-8 hours before fixation.

**FACS**

Isolated cells resuspended in cell sorting buffer (1% BSA, 25 mM HEPES in HBSS) were passed through a 35 μm cell strainer (Corning, #352235) before sorting. $GFP^+$ CFs were sorted using a FACSaria II system (BD Biosciences). Gates were made based on $GFP^-$ controls. Sorted CFs were collected in TRIzol, cell sorting buffer, or complete medium, depending on the downstream experiments.

**Chondrogenesis**

Briefly, $2.5 \times 10^4$ CFs were resuspended in 5 ml of pre-warmed chondrogenic base media within 15 ml centrifuge tubes, followed by centrifugation at 200 x g for 5 minutes at room temperature. The pellet was then resuspended in 0.5 ml of pre-warmed complete chondrogenic differentiation media (ThermoFisher, #A10071-01). To allow gas exchange, the tube caps were loosened, and the tubes were incubated upright at 37° C with 5% CO2. The medium was replaced every 3 days, and the samples were harvested after 14 days of culture. Cell pellets were fixed in 4% PFA for 30 minutes at 4°C, washed three times in 1X PBS, embedded in OCT, and then immediately frozen at -80°C. Cryosections (5 μm) were stained with Alcian Blue (pH 2.5) for 1 hour to evaluate the chondrogenic efficiency.

**Bone marrow derived macrophage (BMDM) isolation and culture**

Bone marrow was flushed out from femur and tibia using a 10 mL syringe filled with ice-cold PBS and fitted with a 26G needle into a 50 mL tube on ice. The suspension was filtered through a 70 μm cell strainer, centrifuged at 500 × g for 10 minutes at 4°C, and the pellet was resuspended in 2 mL of 1× RBC lysis buffer. After 2 minutes on ice, 20 mL of ice-cold PBS was added, followed by centrifugation under the same conditions. The final pellet was resuspended in 6 mL of complete bone marrow medium (DMEM with 10% bovine growth serum and 12.5 ng/mL M-CSF) per mouse. Cells were plated in 12-well plates (1 mL/well) with or without 100 ng/mL PTN and cultured at 37°C with 5% $CO_2$. On day 3, the medium was replaced with fresh medium 5 hours before RNA collection.

**Phagocytosis assay**

Isolated BMDMs were seeded into 96-well plates and cultured for 3 days in complete bone marrow medium with or without 100 ng/mL PTN. Cells were then incubated with pHrodo™ Red Zymosan BioParticles™ Conjugate in the presence of 10% BGS according to the manufacturer's protocol. Images were acquired after 4 hours of incubation using an ECHO Revolve fluorescence microscope.

**Viral packaging**

To generate retroviruses expressing HA-tagged (N-terminal) mouse *Runx1* (NCBI NM_001111023.2) and the human RUNX1 (NCBI NM_001754.5), Platinum-E cells were transfected with MIGR1 (Addgene, #27490) vectors carrying *Runx1*/*RUNX1* cDNA. Transfection was performed using the JetPRIME transfection reagent (Polyplus, #101000015) at a 1:2 DNA to transfection reagent ratio. Retroviral particles were purified following the established ultracentrifugation protocol, using a 25% sucrose cushion prepared in 50 mM sodium phosphate buffer (pH 7.4). The 25% sucrose solution was carefully layered beneath the viral supernatant in an ultracentrifuge tube at a 1:5 ratio (sucrose to supernatant) before centrifugation.

#### Realtime PCR

Briefly, total RNA was isolated using the Direct-zol RNA Microprep Kit (Zymo, #R2063-A) and reverse-transcribed into cDNA using the iScript cDNA Synthesis Kit (Bio-Rad, #1708891). Quantitative PCR was conducted on a QuantStudio™ 3 Real-Time PCR System using SsoAdvanced Universal SYBR Green Supermix (Bio-Rad, #1725274). Gene expression levels were normalized to 18S rRNA, and primer sequences are shown in **Table S1**.

#### Bulk RNAseq

Total RNA was extracted from FACS-sorted $GFP^+$ CFs at different time points (2h, 12h, 1D post-MI) using Qiagen miRNeasy Micro Kit (#217084). cDNA libraries were constructed using the NEBNext single cell/low input RNA library prep kit for Illumina (#E6420). The cDNA libraries were sequenced on the Illumina platform using 150 bp paired-end sequencing. Two biological replicates were included for each time point.

#### Bulk ATACseq

Bulk ATACseq was performed as described previously[4]. Briefly, 10,000 sorted CFs were centrifuged at 1000 × g for 10 minutes, and the supernatant was carefully removed using a 200 µL pipette, leaving approximately 1–2 µL of residual. Cells were resuspended in 8.5 µL of ice-cold 1× lysis buffer and incubated on ice for 20 minutes to allow cytoplasmic lysis. Subsequently, 12.5 µL of 2x Tris-DMF-Tagmentation buffer and 2 µL of Nextera Tn5 transposase (Illumina, #20034198) were added, and the mixture was gently pipetted. The reaction was incubated at 37°C for 30 minutes with shaking, followed by the addition of 0.5 µL of 10% SDS to terminate the reaction. The mixture was gently pipetted again and incubated on ice for 5 minutes to denature Tn5 and release bound DNA. DNA was purified using the QIAGEN MinElute Reaction Cleanup Kit (#28204), and libraries were constructed using Q5® High-Fidelity 2X Master Mix (NEB, #M0492L), followed by purification with SPRI magnetic beads. The ATACseq libraries were sequenced on the Illumina platform using 150 bp paired-end sequencing. Two biological replicates were included for each time point.

#### Bulk CUT&Tag

CUT&Tag was performed following the EpiCypher CUTANA™ CUT&Tag DIY Protocol with modifications. For CFs isolated from hearts of WT mice at different time points, CFs were seeded on 6 cm dishes for 1 hour after FACS and washed once with 2 ml pre-warmed media, followed by the addition of 2 ml pre-warmed media. Light cross-linking was performed by adding 5.72ul 37% formaldehyde (final concentration is 0.1% formaldehyde) while swirling the dish for 1 min and quenching the cross-linking by adding 100ul of 2.5M glycine. CFs were scraped from the plate, transferred to 1.5ml tubes, and centrifuged at 600 g for 4 min at room temperature. 50,000 Cross-linked CFs per reaction were subjected to nuclei preparation and binding nuclei to activated ConA beads (Cell Signaling Technology, #93569) using 8-strip PCR tubes. The unbound supernatant was removed, and bead-bound nuclei were resuspended in 50 µL of cold Antibody150 Buffer (20 mM HEPES pH 7.5; 150 mM NaCl; 0.5 mM Spermidine; 1× EDTA-free Protease inhibitor cocktail; 2 mM EDTA) containing 1:50 diluted primary antibody. Primary antibody incubation was performed on a nutator overnight at 4 °C. The supernatant containing the primary antibody was removed by placing the tube on a magnet stand to clear. After the second wash, 50 µL cold Wash150 Buffer (20 mM HEPES pH 7.5; 150 mM NaCl; 0.5 mM Spermidine; 1× EDTA-free

Protease inhibitor cocktail) containing 1:50 diluted Guinea Pig anti-Rabbit IgG secondary antibody was added to bead-bound nuclei, and nuclei were incubated on nutator at RT for 30 min. Bead-bound nuclei were washed twice for 5 min each in 200 μL cold Wash150 Buffer on the magnet to remove unbound antibodies. After the second wash, 50 μL Wash300 Buffer (20 mM HEPES pH 7.5; 300 mM NaCl; 0.5 mM Spermidine; 1× EDTA-free Protease inhibitor cocktail) containing 2.5 μL CUTANA pAG-Tn5 (20x stock, EpiCypher, #15-1017) was added to each reaction. After gently pipetting, the nuclei were incubated on a rotator for 1 hour at RT. Bead-bound nuclei were washed twice for 5 min each in 200 μL cold Wash300 Buffer with thorough pipetting before being put onto the magnet to remove unbound CUTANA pAG-Tn5. Next, Bead-bound nuclei were resuspended in 50 μL Tagmentation buffer (10 mM MgCl2 in Wash300 Buffer) and incubated at 37 °C for 1 hour in a thermocycler. Beads were resuspended in 50 μL RT TAPS buffer (10 mM TAPS, pH 8.5; 0.2 mM EDTA) by pipetting, and then the supernatant was discarded after placing the tube on the magnet stand to clear. The beads were resuspended by vortexing using 5 μL RT SDS Release Buffer (10 mM TAPS, pH 8.5; 0.1% SDS) to stop tagmentation. The tubes were incubated at 58 °C for 1 hour in a thermocycler. After incubation, 15 μL RT SDS Quench Buffer (0.67% Triton-X 100 in Molecular grade $H_2O$) was added to each reaction to neutralize SDS. To amplify libraries, 2 μL universal i5 and 2 μL barcoded i7 primer (New England Biolabs) and 25 μL NEB Q5® High-Fidelity 2X Master Mix (#M0492L) were added to each reaction. The sample DNA was amplified for 14-18 cycles using CUT&Tag-specific PCR cycling parameters. Post-PCR clean-up was performed by adding 1.3X volume of AMPure XP beads (Beckman Counter, #A63881), incubating for 15 min at RT, washing twice gently in 80% ethanol, and eluting in 25 μL Molecular grade $H_2O$. Cultured CFs were subjected to CUT&Tag following the same protocol described above without fixation. Two biological replicates were included for each time point.

**Bulk CUT&RUN**

CFs isolated from WT C57BL/6J mice were transduced with retroviruses expressing HA-tagged (N-terminal) mouse *Runx1* (NCBI NM_001111023.2) and the human RUNX1 (NCBI NM_001754.5). Two days after transduction, 500,000 CFs per reaction were subjected to CUT&RUN using Epicypher CUT&RUN DIY protocol with modifications. Light cross-linking was performed by adding 5.72ul 37% formaldehyde to 2ml CF suspension in 1X PBS for 2 minutes, followed by quenching by adding 100ul of 2.5M glycine 125mM glycine. Cross-linked CFs were bond to activated ConA beads. The unbound supernatant was removed, and bead-bound CFs were washed with XL Wash Buffer (20 mM HEPES pH 7.5; 150 mM NaCl; 0.5 mM Spermidine; 1× EDTA-free Protease inhibitor cocktail; 1% Triton X-100; 0.05% SDS) twice and then resuspended in 50 μL of cold Antibody Buffer (20 mM HEPES pH 7.5; 150 mM NaCl; 0.5 mM Spermidine; 1× EDTA-free Protease inhibitor cocktail; 1% Triton X-100; 0.05% SDS; 0.01% Digitonin; 2 mM EDTA) containing 0.5 ug primary antibody. Primary antibody incubation was performed on a nutator overnight at 4 °C. The supernatant containing the primary antibody was removed by placing the tube on the magnet stand to clear and the beads were washed two times using XL Digitonin Buffer (20 mM HEPES pH 7.5; 150 mM NaCl; 0.5 mM Spermidine; 1× EDTA-free Protease inhibitor cocktail; 1% Triton X-100; 0.05% SDS; 0.01% Digitonin). After the second wash, 50 μL XL Digitonin Buffer containing 2.5 μL pAG-MNase (20x stock, Epicypher, #15-1016) was added per reaction to bead-bound cells, and cells were incubated on a nutator at RT for 10 min. The supernatant containing pAG-MNase was removed and the beads were washed twice using cold XL Digitonin Buffer. After the second wash, 50 μL XL Digitonin Buffer containing 1 μL 100 mM $CaCl_2$ was added to bead-bound cells per reaction, and cells were incubated on a nutator at 4ºC for 2 hours. After incubation, 33 μL Stop Buffer (20 mM EDTA; 340 mM NaCl; 4 mM EGTA; 50 ug/ml RNase A; 50 ug/ml Glycogen) with an optimized amount of E. coli DNA was added to each reaction. Cells were incubated for 10 min at 37ºC in a thermocycler. After incubation, the tubes were placed on a magnet, and the supernatant containing fragmented CUT&RUN-enriched

DNA was transferred to new 8-strip PCR tubes. For each reaction, 0.8 µL 10% SDS and 1 µL of 20 µg/µL Proteinase K were added, followed by overnight incubation at 55ºC using a thermocycler to reverse cross-links. DNA was purified using QIAGEN MinElute Reaction Cleanup Kit (#28204) and eluted in 20 µL Elution Buffer. For each reaction, 5 ng was used to prepare Illumina NGS libraries using NEBNext® Ultra™ II DNA Library Prep Kit (E7645). Two biological replicates were included for each time point.

**Bulk Hi-C**

Hi-C library was constructed from 700,000 FACS-sorted CFs from uninjured or 7D post-MI hearts using the Arima-HiC kit and Arima Library Prep Module (Arima Genomics, A510008) according to the manufacturer's protocol. The Hi-C libraries were sequenced on the Illumina platform using 150 bp paired-end sequencing. 500-700 million read pairs were obtained for each sample. Two biological replicates from each time point were analyzed.

**Enzymatic methyl-seq**

Enzymatic methyl-seq libraries were constructed using the NEBNext® Enzymatic Methyl-seq Kit (New England Biolabs, #E7120). Briefly, genomic DNA was isolated from sorted CFs using the Quick-DNA Microprep Kit (Zymo Research, #D3020) and mixed with control DNA. Combined DNA was fragmented to an average size of ~250 bp using a Covaris M220 Focused-ultrasonicator (Covaris, Woburn, MA, USA). After fragmentation, the sheared DNA was ligated to adaptors. In the subsequent enzymatic conversion steps, 5-methylcytosines and 5-hydroxymethylcytosines were protected by TET2 and Oxidation Enhancer and APOBEC was applied to convert unmodified cytosines to uracils. Finally, PCR amplification and purification were performed to enrich the libraries. The libraries were sequenced on the Illumina platform using 150 bp paired-end sequencing. Two biological replicates from each time point were analyzed.

**Single-nucleus multiome (RNAseq+ATACseq)**

Three hearts of each treatment group were combined and minced in ice-cold isotonic lysis buffer (10 mM Tris-HCl pH 7.4, 150 mM NaCl, 3 mM $MgCl_2$, 1% BSA; no detergent or DTT). Minced tissue was allowed to settle, and excess buffer was removed. Tissue fragments were transferred to a Dounce homogenizer (Electron Microscopy Sciences, #64790-01) in isotonic lysis buffer containing 1 mM DTT, homogenized with 10 strokes using the loose pestle, filtered through a 40 µm cell strainer, and centrifuged at 500 × g for 5 min at 4 °C. Pellets were resuspended in isotonic lysis buffer with 1% BSA; an aliquot was stained with DAPI to assess nuclear integrity, and the remaining suspension was stained with 7-AAD and the 7-AAD$^+$ fraction was sorted into tubes prefilled with isotonic lysis buffer + 1% BSA. Sorted samples were centrifuged (500 × g, 5 min, 4 °C), resuspended in 0.1× hypotonic lysis buffer (10 mM Tris-HCl pH 7.4, 10 mM NaCl, 3 mM $MgCl_2$, 0.01% NP-40, 1% BSA, 1 mM DTT), incubated on ice for 2 min, diluted with hypotonic lysis dilution buffer (without NP-40), and centrifuged again. The final nuclear pellets were resuspended in freshly prepared 10x Genomics diluted nuclei buffer (1× nuclei buffer from 20× stock, 1 mM DTT, 1 U/µl RNase inhibitor), maintained on ice, and nuclei were counted by DAPI staining using an ECHO microscope. Isolated nuclei were used to construct single-nucleus Multiomic libraries using the 10x Genomics Chromium Next GEM Single Cell Multiome ATAC + Gene Expression kit (#PN-1000283). Twenty thousand nuclei were used for each reaction. Libraries were sequenced on an Illumina platform using 150 bp paired-end sequencing.

**Single-cell RNAseq**

Cells isolated from 3 mouse hearts per treatment group were combined and stained with the Live/Dead kit (Thermo Fisher, #L3224) and Hoechst dye (AAT Bioquest, #17537). Cells that were negative for EthD-1 and positive for calcein or Hoechst were collected, centrifuged, and adjusted to 1,000-2,000 cells/μL. For each sample, 20,000 cells were used to construct library using Chromium Single Cell 3′ Reagent Kits v3 (10x Genomics). Libraries were sequenced on an Illumina platform using 150 bp paired-end sequencing.

**Bulk RNAseq analysis**

To remove low-quality base calls, adapters, and short reads, TrimGalore (v0.6.7) was used with the following options: *--quality 20 --stringency 1 --clip_R1 9 --clip_R2 9*, with all other parameters set to default. After quality assessment using FastQC, clean reads were aligned to the mouse genome (mm10) using STAR (v2.7.10b) in two-pass mode. For both passes, STAR was executed following ENCODE project recommendations, largely using default settings with a few exceptions (e.g., alignSJoverhangMin=8, alignIntronMin=20). Gene- and transcript-level expression was then quantified using RSEM (v1.3.3) to obtain raw read counts and transcripts per million (TPM). The R package DESeq2 (https://github.com/thelovelab/DESeq2) was used to perform differential gene expression analysis with accounting for batch effects. A gene was considered differentially expressed if the absolute shrunken $\log_2$ fold change (LFC) in TPM exceeded 0.585 and the adjusted p-value (padj) was below 0.05.

**Bulk ATACseq analysis**

As in bulk RNAseq processing, sequencing reads from all samples were first subjected to adapter and low-quality base calls trimming and removal of short fragments using TrimGalore (v0.6.7), followed by quality assessment with FastQC. Reads were then aligned to the mouse reference genome (mm10) using Bowtie2 (v2.4.1) with the options: --very-sensitive -X 2000 --no-mixed --no-discordant. Only uniquely mapped reads were retained for downstream analysis, and PCR duplicates as well as reads mapping to mitochondrial DNA were removed.

Peaks were called separately for each sample using MACS2 (v2.2.8) with the options: --keep-dup all --nolambda --nomodel. For each biological condition (e.g., uninjured hearts), peaks from individual samples were merged using bedtools to obtain condition-specific open chromatin regions. To enable differential accessibility analysis, peaks from all samples included in a comparison were merged using bedtools to generate a unified peak set, and accessibility for each peak in each sample was quantified as fragments per million mapped reads (FPM).

To quantify chromatin accessibility genome-wide, the mouse genome was segmented into consecutive non-overlapping 300-bp bins, and FPM values were computed per bin for each sample. Promoter accessibility was assessed by calculating FPM across promoter regions, defined as ±500 bp around the transcription start site (TSS) of each annotated gene.

Genomic feature annotations, including TSSs, transcription end sites (TESs), exons, introns, and among others, were obtained from the UCSC Genome Browser. Intergenic regions were defined as the genomic intervals upstream of the first TSS and downstream of the last TES on each chromosome, as well as the regions between the TES of one gene and the TSS of the next. Peaks not overlapping annotated promoter regions were classified as distal peaks.

To evaluate enrichment of genomic features among ATACseq peaks, a set of random peaks matched in length to the observed peaks was generated. Enrichment was calculated as the ratio (or log ratio) of the number of ATACseq peaks overlapping each genomic feature to the number of random peaks overlapping the same feature. TF motif enrichment in peaks was assessed using the findMotifsGenome.pl command from HOMER, and motif locations within peaks were identified using the findMotifsGenome.pl command.

Differential accessibility analysis of both peak regions (i.e., those included in the unified set described above) and promoters across biological conditions was performed using DESeq2 with accounting for batch effects. A peak region or promoter was considered with differential accessible if the absolute shrunken LFC in FPM exceeded 0.585 and the padj was below 0.05.

**Bulk CUT&Tag analysis**

The data processing and analyses were performed as described for ATACseq dataset, with the exception that MACS2 peak calling was executed with the options: --keep-dup all --nolambda.

**Bulk Cut&Run analysis**

Data processing and analysis followed the same workflow as for the CUT&Tag datasets.

**Hi-C data analysis**

Sequencing data from individual samples were processed using the Juicer pipeline (v1.6, https://github.com/aidenlab/juicer) with the mouse genome (mm10) as the reference. A restriction site file specific to the mm10 genome was first generated using the generate_site_positions.py script provided in Juicer. Sequencing reads were then processed with juicer.sh using the generated restriction site file to produce .hic files containing contact matrices. These .hic files were subsequently analyzed with the Arrowhead algorithm to identify contact domains and with the HICCUPS algorithm to detect chromatin loops at multiple resolutions (5 kb, 10 kb, and 25 kb).

With .hic files from Juicer, differential compartment, subcompartment, and chromatin loop analyses were performed using dcHiC (v1.0.0, https://github.com/ay-lab/dcHiC). The .hic files were converted into dense contact matrices and binned at 100-kb resolution across the mm10 genome. For each chromosome, dcHiC computed the observed/expected (O/E) contact matrices and applied principal component analysis (PCA) to derive the first eigenvector (PC1), representing large-scale chromatin compartmentalization. Positive PC1 values correspond to transcriptionally active A compartments, whereas negative values correspond to transcriptionally inactive B compartments.

PC1 vectors were z-score normalized and compared across biological conditions to identify differential chromatin compartments. Genomic bins with $|\Delta PC1| \geq 0.4$ and a false discovery rate (FDR) $< 0.05$ were considered significantly altered. Each bin was further classified into one of four subcompartments: strong A (sA), weak A (wA), weak B (wB), and strong B (sB). The sA subcompartment corresponds to the most transcriptionally active, gene-dense regions with high contact frequency; wA represents moderately active chromatin; wB reflects partially condensed chromatin with low transcriptional activity; and sB denotes strongly repressed, lamina-associated heterochromatin.

To categorize structural transitions between biological conditions (e.g., from uninjured to post-MI), bins were compared based on changes in compartment sign and the magnitude of their PC1 (|PC1|). Cross-compartment switches were defined as A→B transitions, indicating a shift from active to repressed states and B→A transitions, indicating a shift from repressed to active states. Within-compartment shifts were classified according to changes in |PC1|: bins that remained in compartment A but showed increased |PC1| were designated wA→sA (weak-to-strong A), whereas those with decreased |PC1| were designated sA→wA (strong-to-weak A). Likewise, bins remaining in compartment B with increased |PC1| were classified as wB→sB (weak-to-strong B), and those with decreased |PC1| as sB→wB (strong-to-weak B). Bins with invalid compartment scores in either condition were excluded from above classification.

Long-range interactions (i.e., loops) were identified from contact matrices using FitHiC (v2.0.8, https://github.com/ay-lab/fithic). FitHiC models the contact probability as a function of genomic distance and computes *p*-values for each interaction while accounting for distance-dependent background biases. Significant loops were called at the 100kb resolution with an FDR < 0.05. Differential loop strength between conditions was assessed based on differences in contact enrichment. Loop anchors were annotated to nearby genes and regulatory elements to identify promoter–enhancer interactions altered in response to MI.

All analyses were conducted using the R implementation of dcHiC, and the visualization of compartments, subcompartments, and loops was carried out using Juicebox in combination with matplotlib and seaborn in Python.

**Enzymatic methyl-seq analysis**

After adapter and low-quality read removal using TrimGalore (v0.6.7) and quality assessment with FastQC, as performed for RNAseq data processing, Bismark (v0.22.3) was used to align reads to the mouse genome (mm10) with the following parameters: --un --non_directional --score_min L,0,-0.6. Duplicate alignments were then removed using the deduplicate_bismark command, followed by quantification of methylation levels at individual CpG sites with the bismark_methylation_extractor command. The methylation level at each CpG site was defined as the percentage of methylated reads among all reads overlapping that site. For downstream analyses, only CpG sites covered by at least three reads were retained.

Promoter methylation levels were calculated as the average methylation level across CpG sites within each promoter region. To assess genome-wide methylation patterns, the genome was partitioned into non-overlapping 300 bp bins, and the methylation level of each bin was computed as the mean methylation level across all CpG sites within it.

**Super-enhancer Identification**

Super-enhancer was identified using the Rank Ordering of Super-Enhancers (ROSE, v1.3.2; https://github.com/stjude/ROSE) pipeline with default parameters. In brief, ATACseq BAM files, UCSC gene annotation files, and peak files were provided as input. The pipeline ultimately generated BED files containing stitched enhancer regions classified as super-enhancers.

**Bulk sequencing data integration**

To categorize genes by expression level, we computed a Z-score for each gene in each sample by standardizing TPM values across all genes within the sample to a mean of 0 and variance of 1. Genes with Z-scores consistently below (or above) zero across all samples were designated as having constitutively low (or high) expression, while the remaining genes were classified as dynamically expressed. The classification of genes by individual promoter epigenetic states including accessibility and histone modifications (i.e., H3K27ac, H3K27me3, and H3K4me3), i.e., constitutively low, constitutively high, and dynamic, was done similarly.

Promoter epigenetic states and gene expression levels for each biological condition (e.g., uninjured hearts and hearts at 3D post-MI) were obtained by averaging values across samples within the same condition. To assess the relationship between promoter epigenetic states and gene expression, Pearson correlation coefficients were calculated across all biological conditions, and promoters showing strong correlations ($cor \geqslant 0.5$) were identified. Similar correlation was also computed between distal peak epigenetic states and gene expression with peaks linked to their closest genes by linear distance. Heatmaps of gene expression and promoter or distal peak epigenetic states were generated using the ComplexHeatmap R package.

To further assess the association among distinct epigenetic modalities and gene expression, we first applied the same approach described above to compute correlations between each pair of epigenetic modalities at gene promoters across biological conditions. We additionally calculated correlations among epigenetic modalities together with gene expression across gene promoters within individual biological conditions. All analyses were restricted to promoters of genes that exhibited differential expression across the corresponding conditions. Heatmaps and violin plots were generated using Python packages: Seaborn and Matplotlib.

**Single-nucleus multiome analysis**

To obtain the RNA-ATAC multiome matrix for downstream analysis, raw FASTQ files were firstly aligned to the mm10 reference genome (https://www.10xgenomics.com/support/software/cell-ranger-arc/downloads) using Cell Ranger ARC (v2.0.2, 10x Genomics, Pleasanton, CA, USA). After Cell Ranger ARC alignment, quality control was performed using a multi-step pipeline. Ambient RNA was removed with CellBender (v0.2.0; FPR=0.01, 150 epochs, https://github.com/broadinstitute/CellBender). RNA quality control used universal minimum thresholds (UMI: ≥300; genes: ≥100) and cluster-specific dynamic maximum thresholds for UMI count, gene count, and mitochondrial UMI fraction that were calculated using 75th percentile plus 1.5× interquartile range (IQR) for each cluster. Doublet detection was performed separately: RNA using DoubletFinder (pK optimized via parameter sweeping, nExp = 0.000008 × cell count, homotypic/heterotypic classification, https://github.com/chris-mcginnis-ucsf/DoubletFinder), and ATAC using ArchR (https://github.com/GreenleafLab/ArchR). ATAC quality control filtered cells by TSS enrichment (≥1), nucleosome signal (≤2), and fragment counts (minimum: 2,000), with cluster-specific maximum fragment count thresholds calculated dynamically using the 75th percentile plus 1.5× IQR method. Final high-quality nuclei that passed all RNA and ATAC filters, and dual-modality doublet removal (excluded if labeled as doublet in either modality) were used for subsequent analysis.

After getting the high-quality matrix, Seurat (v5.2.1, https://satijalab.org/seurat/) and Signac (v1.14.0, https://stuartlab.org/signac/) packages in R (v4.4.2) were used for cluster analysis following the standard pipeline. For the RNA modality, data were first normalized using SCTransform (https://satijalab.org/seurat/articles/sctransform_vignette.html) or LogNormalize. Then, dimensionality reduction was performed using Uniform Manifold Approximation and Projection (UMAP), and clustering was conducted using the Leiden algorithm, while all other parameters were kept at their defaults. For the ATAC modality, peaks were first called using MACS2. The resulting peak-by-cell matrix was normalized, and latent semantic indexing (LSI) was performed using dimensions 2–50 to capture major sources of chromatin accessibility variation. Subsequently, a shared nearest-neighbor (SNN) graph was constructed based on the LSI embeddings to enable downstream clustering and visualization. For the multi-modal analysis, FindMultiModalNeighbors was first applied to construct a joint neighbor graph (WNN), followed by Leiden clustering with the resolution specified in the Results section. UMAP embeddings were then generated on the WNN graph using RunUMAP with default parameters. Finally, differentially expressed genes between clusters were identified using FindAllMarkers.

Gene activity scores were computed from ATACseq data to estimate transcriptional potential from chromatin accessibility. Scores were calculated using the GeneActivity() function (Signac v1.9.0) with default parameters, aggregating ATACseq fragment counts within gene bodies and promoter regions. Scores were normalised using SCTransform (Seurat v5.2.1) to create the ACTIVITY_SCT assay. Downstream analysis was performed on the ACTIVITY_SCT assay.

To identify co-accessibility predicted interactions among loci, preprocessed scATACseq Seurat objects were analyzed in R using Signac (v1.14.0, https://stuartlab.org/signac/) and Seurat (v5.2.1, https://satijalab.org/seurat/). Co-accessible peak–peak interactions were inferred using Cicero (Monocle, https://stuartlab.org/signac/articles/cicero) and filtered to retain links with co-accessibility score > 0.10. Aggregated accessibility profiles were visualized by cluster and sample together with gene annotations, peak calls, and co-accessibility arcs. External RUNX1/Runx1 Cut&Run BigWig tracks were overlaid as smoothed coverage tracks.

**Gene regulatory network construction**

The SCENIC+ (https://github.com/aertslab/scenicplus) pipeline was employed to reconstruct gene regulatory networks (GRNs) from single-nucleus RNAseq and ATACseq datasets. The major steps and our specific adaptations for the reconstruction are summarized below; for a detailed description and discussion, please refer to the SCENIC+ publication.

To generate a set of consensus peaks, pseudobulk ATACseq profiles were first created for each cell type by aggregating aligned fragments from cells of the same type. MACS2 was then applied to each pseudobulk profile to identify peaks, which were subsequently merged across all profiles to define the final set of consensus peaks, as described in SCENIC+. The accessibility of these consensus peaks in individual cells was quantified by counting the number of overlapping fragments, yielding a count matrix.

A Markov chain Monte Carlo (MCMC)-based topic finding algorithm was applied to the count matrix above to identify "hot topics," producing two matrices that capture the relationships between cells and topics (cell–topic) and between topics and regions (topic–region). The product of these two matrices was then used to impute dropouts in the ATACseq profiles of individual cells. To associate TFs with genomic regions, motif enrichment analysis using pycisTarget was performed on topic-associated regions—identified by binarizing the topic–region matrix—and on regions showing differential accessibility across cell types (padj < 0.05 and LFC > 0.5). Two binarization methods were used: "otsu" [reference] and "ntop" with N = 3,000. Default parameters were applied when running pycisTarget. The ranking- and score-based databases required by pycisTarget were obtained from https://resources.aertslab.org/cistarget/databases/mus_musculus/mm10/screen/mc_v10_clust/region_based/.

To link genomic regions to genes, which is to identify gene enhancers, we incorporated both Hi-C and H3K27ac histone modification profiles. By default, SCENIC+ defines the candidate enhancer search space for a given gene as the genomic region spanning 150 kb upstream of its transcription start site and 150 kb downstream of its end. In our study, when Hi-C profiles were considered, this search space was expanded to include the boundaries of contact domains overlapping with gene promoters. Contact domains identified from all four Hi-C datasets (two from uninjured hearts and two from hearts at 7D post-MI) were merged and applied for the search space extension.

For predicting which consensus peaks within the search space act as gene enhancers, SCENIC+ by default trains gradient boosting machines (GBMs) to predict gene expression based on the accessibility of the corresponding peaks. The resulting GBM-derived importance scores were then used to quantify the strength of regulatory relationships between regions and genes. In addition to this default approach, we also computed Pearson correlations between H3K27ac signal intensity at peaks and gene expression levels, and used these correlations as alternative importance scores for enhancer prediction.

With the obtained TF-to-region and region-to-gene relationships, we followed the default procedures of the SCENIC+ pipeline to construct eRegulons, and subsequently quantified their activities in individual cells using the area under the curve (AUC) metric. For gene expression data, we explored the use of both log-transformed and SCTransform-normalized RNAseq data.

#### Single-cell RNAseq analysis

Single-cell RNAseq data processing and analysis were performed following the same procedure described above for the RNAseq modality of single-nucleus multiome data except that a minimum UMI threshold of 500 was used.

#### Cell-cell communication analysis

To investigate the changes in cell–cell communication between different genotypes, CellChat (v2.1.0; https://github.com/jinworks/CellChat) was used following the tutorial "Comparison analysis of multiple datasets using CellChat," with default parameters.

#### Trajectory analysis using MOSCOT

To model temporal transitions of CFs following MI, we applied MOSCOT (v0.3.3, https://github.com/theislab/moscot), an optimal-transport–based framework for mapping cell populations across time-resolved single-cell datasets. MOSCOT formulates this task as a series of optimal transport (OT) couplings between populations sampled at successive time points, with transport cost reflecting transcriptomic similarity in a low-dimensional embedding space.

To generate the required embeddings, we first performed PCA on single-nucleus RNAseq profiles from multiome datasets, and used the resulting low-dimensional representations as inputs to MOSCOT. A TemporalProblem was instantiated using the temporal ordering of samples—uninjured, 2H, 1D, 3D, 7D, and 4D post-MI. OT problems were defined and solved sequentially between each pair of adjacent time points using entropic regularization ($\varepsilon = 1\times10^{-3}$) to ensure stable, smooth couplings. These couplings were subsequently used to infer CF state transitions across the injury–healing process and to visualize temporal progression in the embedding space. All analyses were conducted using MOSCOT's Python implementation.

#### Pseudotime analysis using CellRank2

To reconstruct CF state transitions at individual time points, we applied CellRank2 (v2.0, https://github.com/theislab/cellrank2_reproducibility), a framework for unified fate mapping across multimodal single-cell data. The scRNAseq profiles of uninjured and post-MI hearts were first embedded in a low-dimensional space using PCA. We then used Palantir (v1.1) to generate an initial continuous pseudotime trajectory. The early CF population, i.e., quiescent CFs from uninjured hearts was manually specified as the starting state, allowing Palantir to infer a biologically constrained progression toward activated, proliferative, and differentiated CF states. The resulting Palantir pseudotime values were subsequently imported into CellRank2, which incorporates pseudotime as an external "view" to construct a transition kernel describing the probability of movement between cell states. Using this kernel, CellRank2 computed a macrostate decomposition, grouping transcriptionally similar cells into discrete dynamic states. From the macrostates, CellRank2 automatically identified initial, intermediate, and terminal states based on their position in the transition graph and their probability of absorbing incoming trajectories. To visualize the inferred transitions, lineage probabilities were computed using CellRank2's lineage projection model and mapped onto UMAP embeddings. This enabled visualization of the dominant differentiation routes from early CFs toward multiple terminal fates. All analyses were performed using the standard CellRank2 pipeline.

### Statistics

Unless otherwise specified, data are presented as mean ± SD with all data points shown in bar graphs or mean with whiskers showing Min to Max and all data points shown in box graphs. Statistical analyses were performed using GraphPad Prism 10 (GraphPad Software, Inc.). For comparisons between two groups, a two-tailed t-test was applied. When comparing more than two groups, one-way ANOVA followed by Tukey's post hoc test was used to assess statistical significance. Log-rank (Mantel-Cox) test was used to determine the significance of the difference between the survival rates of $Pdgfra^{CreERT2};R26^{GFP}$, and $Pdgfra^{CreERT2};Runx1^{fl/fl};R26^{GFP}$ mice. Significance levels are as follows: *, $p < 0.05$; **, $p < 0.01$; ***, $p < 0.001$; ****, $p < 0.0001$.

**Supplemental table**

Table S1. Sequences of primers used in realtime PCR.

| Target genes | Forward primer (5'-3') | Reverse primer (5'-3') |
|---|---|---|
| *Runx1* | ACTCGGCAGAACTGAGAAATGCTAC | GTGGCGGATTTGTAAAGACGGTG |
| *18S* | GTAACCCGTTGAACCCCATT | CCATCCAATCGGTAGTAGCG |
| *Comp* | CCTGCGACGACGACATAGATGG | TGGTCTGGGTTATCTTTCTGGGGA |
| *Cemip* | TGATGGGAGTCGAGGTCAC | GAGCACTATGGAATTGTCAGGG |
| *Ptn* | CAGAAACCTCGCCCGCACTTTG | CTGCTGATATTGCTGGGACGACA |
| *Cxcl1* | TTGACCCTGAAGCTCCCTTGGTT | GACAGGTGCCATCAGAGCAGTC |
| *Cxcl10* | ATTTTCTGCCTCATCCTGCTGGG | CTATGGCCCTCATTCTCACTGGC |
| *Ccl4* | TTCCTGCTGTTTCTCTTACACCT | CTGTCTGCCTCTTTTGGTCAG |
| *Sdc4* | CCAGGGCAGCAACATCTTTGAG | AGCAGGATCAGGAAAACGGCAAA |
| *Tnf* | GTGATCGGTCCCCAAAGGGATG | ATGATCTGAGTGTGAGGGTCTGG |
| *Cdh2* | ACTTGCCAGAAAACTCCAGAGGAC | GTGACGCTGTATCTCAGGGAAAGG |
| *Lox* | GTAACTGCAAACTGCCACGTCCT | AGCGGAGAAGGGGACAAAGCTG |

### SUPPLEMENTAL FIGURES AND FIGURE LEGENDS

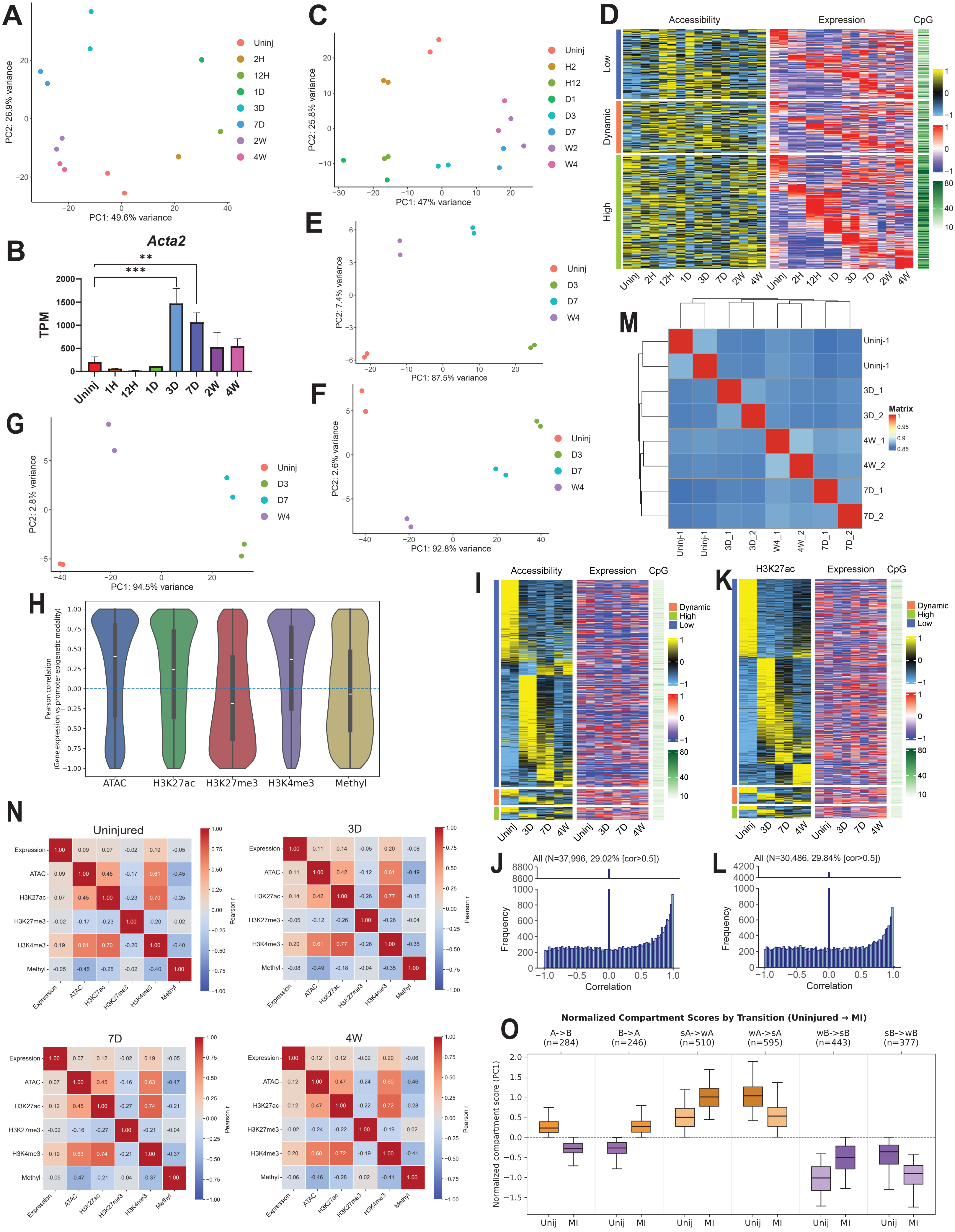

A
PC1: 49.6% variance
PC2: 26.9% variance
Uninj
2H
12H
1D
3D
7D
2W
4W
B
Acta2
TPM
**
***
Uninj
1H
12H
1D
3D
7D
2W
4W
C
PC1: 47% variance
PC2: 25.8% variance
Uninj
H2
H12
D1
D3
D7
W2
W4
D
Accessibility
Expression
CpG
Low
Dynamic
High
E
PC1: 87.5% variance
PC2: 7.4% variance
Uninj
D3
D7
W4
F
PC1: 92.8% variance
PC2: 2.6% variance
G
PC1: 94.5% variance
PC2: 2.8% variance
M
Uninj-1
3D_1
3D_2
4W_1
4W_2
7D_1
7D_2
Matrix
H
Pearson correlation
(Gene expression vs promoter epigenetic modality)
ATAC
H3K27ac
H3K27me3
H3K4me3
Methyl
I
Accessibility
Expression
CpG
Dynamic
High
Low
K
H3K27ac
Expression
CpG
J
All (N=37,996, 29.02% [cor>0.5])
Frequency
Correlation
L
All (N=30,486, 29.84% [cor>0.5])
Frequency
Correlation
N
Uninjured
3D
7D
4W
Expression
ATAC
H3K27ac
H3K27me3
H3K4me3
Methyl
Pearson r
O
Normalized Compartment Scores by Transition (Uninjured → MI)
A->B (n=284)
B->A (n=246)
sA->wA (n=510)
wA->sA (n=595)
wB->sB (n=443)
sB->wB (n=377)
Normalized compartment score (PC1)
Unij
MI

**Figure S1. Bulk multiomic analysis reveals dynamic gene expression and epigenetic regulations in CFs after MI.** (A) A PCA graph shows clustering of bulk RNAseq data from CFs collected from uninjured hearts and at indicated post-MI time points. (B) A bar graph shows *Acta2* expression in CFs across time points by bulk RNAseq. (C) A PCA graph shows clustering of bulk ATACseq data from CFs collected from uninjured hearts and at indicated post-MI time points. (D) A dual-modality heatmap illustrates correlations between DEG expression and promoter accessibility across time points. Promoter CpG densities are also shown. (E-G) PCA graphs show clustering of H3K4me3 (E), H3K27ac (F), and H3K27me3 (G) Cut&Tag data from CFs collected from uninjured hearts and at indicated post-MI time points. (H) A violin plot shows Pearson correlations between gene expression and promoter epigenetic signals for DEGs identified in CFs from uninjured hearts and at 3D, 7D, and 4W post-MI. White bars indicate medians. (I-J) A heatmap (I) and a histogram (J) show correlations between differential distal ATACseq peaks and proximal gene expression across uninjured and post-MI CFs. In heatmaps, Genes are classified into high, low, and dynamic expression groups. CpG densities in ATACseq peak regions are also shown. (K-L) A heatmap (K) and a histogram (L) show correlations between differential distal H3K27ac Cut&Tag peaks and proximal gene expression across uninjured and post-MI CFs. In heatmaps, Genes are classified into high, low, and dynamic expression groups. CpG densities in ATACseq peak regions are also shown. (M) A heatmap shows Pearson correlations among CF EM-seq data from uninjured hearts and at 3D, 7D, and 4W post-MI. (N) Heatmaps show Pearson correlations among indicated modalities across all genes at indicated individual time points. (O) A box plot shows differential compartment transitions between CFs from uninjured hearts and at 7D post-MI (wA: weak A; sA: strong A; wB: weak B; sB: strong B).

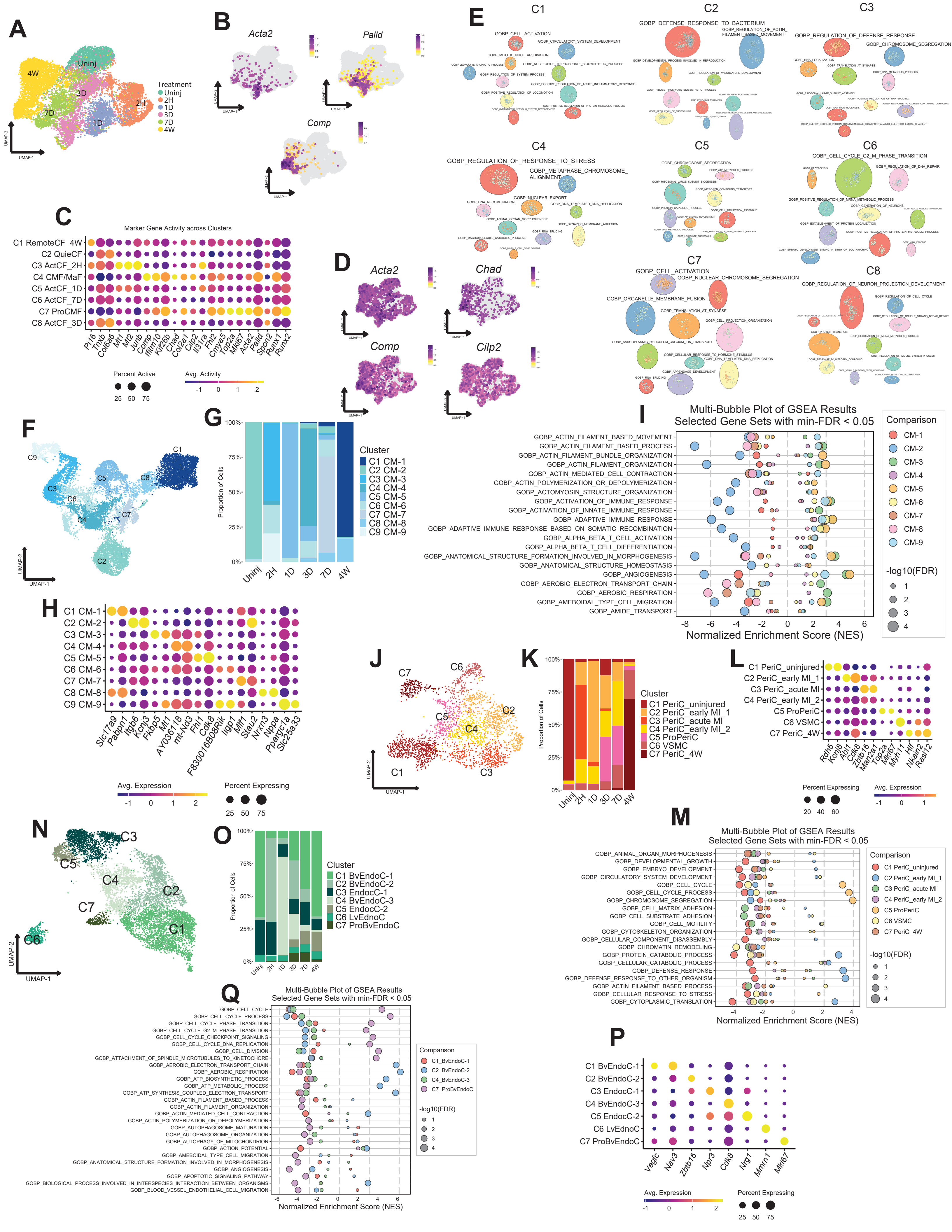

**Figure S2. snMultiomic analysis reveals dynamic gene expression and chromatin accessibility in cardiac cell types after MI.** (A) A multi-modality joint UMAP graph shows CFs from different time points. (B) Feature plots show the expression of selected genes in CFs. (C) A dot plot shows the gene activity score of representative marker genes used for CF subtype annotation. (D) Feature plots show the gene activity score of selected genes in CFs. (E) Graphs show the top 10 GOBP term clusters enriched in individual CF clusters identified by GSEA. (F-G) A multi-modality joint UMAP graph (F) and a bar graph (G) show different CM subtypes identified in mouse hearts without injury and at post-MI time points. (H) A dot plot shows the expression of representative marker genes used for CM subtype annotation. (I) A multi-bubble plot illustrates differential enrichment of selected GOBP terms in CM subtypes identified by GSEA. (J-K) A joint UMAP graph (J) and a bar graph (K) showing MuC subtypes identified across time points. (L) A dot plot shows the expression of representative marker genes used for MuC subtype annotation. (M) A multi-bubble plot shows differential enrichment of selected GOBP terms in MuC subtypes identified by GSEA. (N-O) A multi-modality joint UMAP graph (N) and a bar graph (O) show different EndoC subtypes identified time points. (P) A dot plot shows the expression of representative marker genes used for EndoC subtype annotation. (Q) A multi-bubble plot illustrates the differential enrichment of selected GOBP terms in EndoC subtypes identified by GSEA.

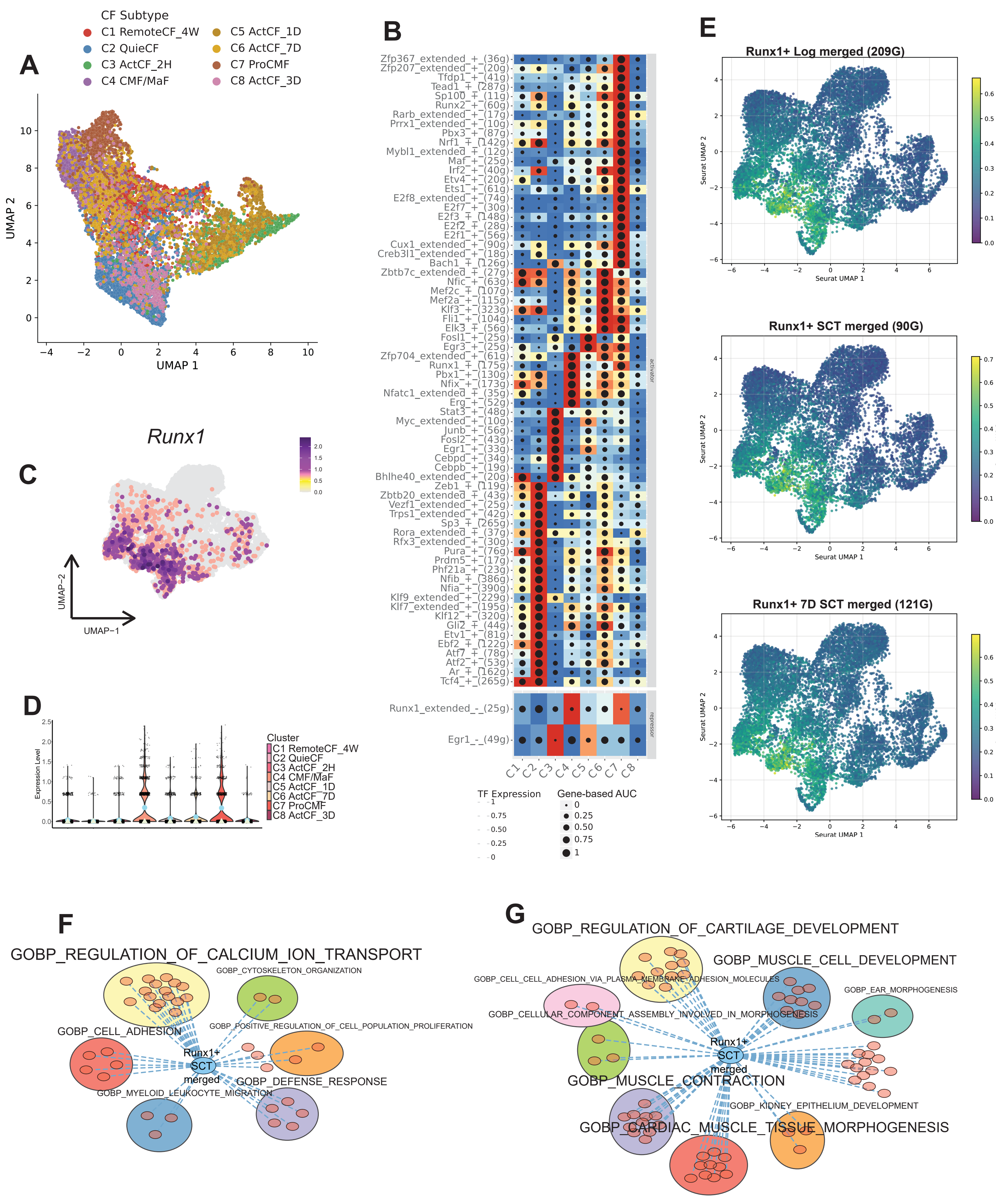

A
CF Subtype
C1 RemoteCF_4W
C2 QuieCF
C3 ActCF_2H
C4 CMF/MaF
C5 ActCF_1D
C6 ActCF_7D
C7 ProCMF
C8 ActCF_3D
UMAP 1
UMAP 2
B
activator
repressor
Runx1_extended_-_(25g)
Egr1_-_(49g)
TF Expression
Gene-based AUC
C
Runx1
UMAP−1
UMAP−2
D
Expression Level
Cluster
E
Runx1+ Log merged (209G)
Runx1+ SCT merged (90G)
Runx1+ 7D SCT merged (121G)
Seurat UMAP 1
Seurat UMAP 2
Custom AUC Score
F
GOBP_REGULATION_OF_CALCIUM_ION_TRANSPORT
GOBP_CYTOSKELETON_ORGANIZATION
GOBP_CELL_ADHESION
GOBP_POSITIVE_REGULATION_OF_CELL_POPULATION_PROLIFERATION
Runx1+ SCT merged
GOBP_DEFENSE_RESPONSE
GOBP_MYELOID_LEUKOCYTE_MIGRATION
G
GOBP_REGULATION_OF_CARTILAGE_DEVELOPMENT
GOBP_MUSCLE_CELL_DEVELOPMENT
GOBP_CELL_CELL_ADHESION_VIA_PLASMA_MEMBRANE_ADHESION_MOLECULES
GOBP_EAR_MORPHOGENESIS
GOBP_CELLULAR_COMPONENT_ASSEMBLY_INVOLVED_IN_MORPHOGENESIS
Runx1+ SCT merged
GOBP_MUSCLE_CONTRACTION
GOBP_KIDNEY_EPITHELIUM_DEVELOPMENT
GOBP_CARDIAC_MUSCLE_TISSUE_MORPHOGENESIS

**Figure S3. Construction of GRNs governing CF gene expression after MI.** (A) A UMAP graph of CFs based on regulon activity identified by SCENIC+ using “sctransform”-normalized data. (B) A heatmap dot plot shows the gene-based regulon AUC scores and corresponding TF expression for the most differentially active regulons across CF subtypes identified using “sctransform”-normalized data. (C-D) A feature plot (C) and a violin plot (D) show *Runx1* expression in different CF clusters. Triangles and dots in the violin plot indicate median and mean, respectively. (E) Feature plots display gene-based AUC scores for indicated versions of the Runx1+ regulon in CFs. (F) A graph shows overlap between “Runx1+ SCT merged” regulon and core enrichment genes in GOBP terms enriched in C7 ProCMF. (G) A graphs shows overlap between “Runx1+ SCT merged” regulon and core enrichment genes in GOBP terms enriched in C4 CMF/MaF.

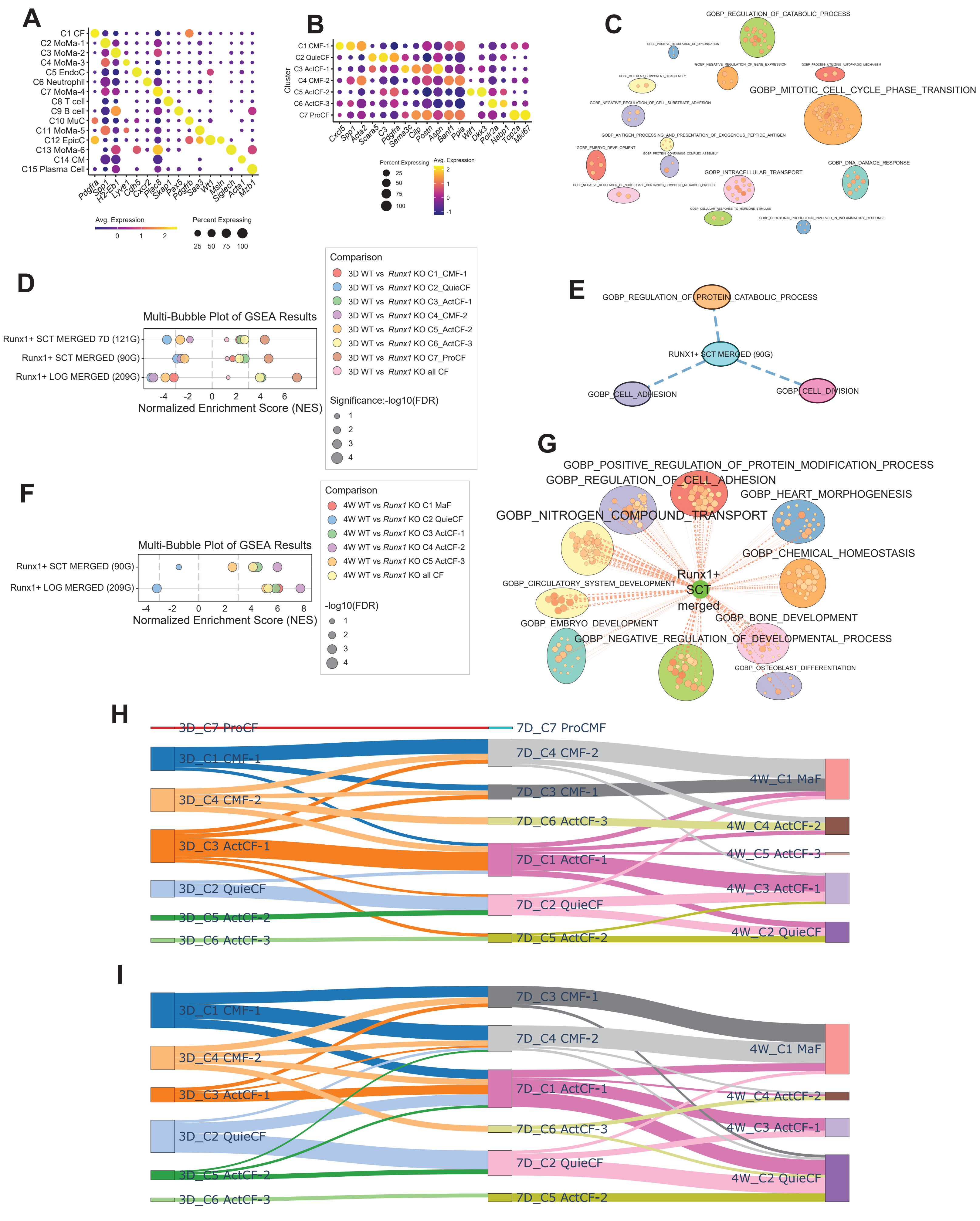

A
C1 CF
C2 MoMa-1
C3 MoMa-2
C4 MoMa-3
C5 EndoC
C6 Neutrophil
C7 MoMa-4
C8 T cell
C9 B cell
C10 MuC
C11 MoMa-5
C12 EpicC
C13 MoMa-6
C14 CM
C15 Plasma Cell
Pdgfra Spp1 H2-Eb1 Lyve1 Cdh5 Cxcr2 Plac8 Skap1 Pax5 Pdgfrb Saa3 Wt1 Msln Siglech Acta1 Mzb1
Avg. Expression
0 1 2
Percent Expressing
25 50 75 100
B
C1 CMF-1
C2 QuieCF
C3 ActCF-1
C4 CMF-2
C5 ActCF-2
C6 ActCF-3
C7 ProCF
Cluster
Cxcl5 Spp1 Acta2 Scara5 C3 Pdgfra Sema3c Cilp Postn Aspn Banf1 Ppia Wif1 Dkk3 Polr2a Nabp1 Top2a Mki67
Percent Expressing
25 50 75 100
Avg. Expression
2 1 0 -1
C
GOBP_REGULATION_OF_CATABOLIC_PROCESS
GOBP_MITOTIC_CELL_CYCLE_PHASE_TRANSITION
GOBP_INTRACELLULAR_TRANSPORT
GOBP_DNA_DAMAGE_RESPONSE
D
Multi-Bubble Plot of GSEA Results
Runx1+ SCT MERGED 7D (121G)
Runx1+ SCT MERGED (90G)
Runx1+ LOG MERGED (209G)
-4 -2 0 2 4 6
Normalized Enrichment Score (NES)
Comparison
3D WT vs Runx1 KO C1_CMF-1
3D WT vs Runx1 KO C2_QuieCF
3D WT vs Runx1 KO C3_ActCF-1
3D WT vs Runx1 KO C4_CMF-2
3D WT vs Runx1 KO C5_ActCF-2
3D WT vs Runx1 KO C6_ActCF-3
3D WT vs Runx1 KO C7_ProCF
3D WT vs Runx1 KO all CF
Significance:-log10(FDR)
1
2
3
4
E
GOBP_REGULATION_OF_PROTEIN_CATABOLIC_PROCESS
RUNX1+ SCT MERGED (90G)
GOBP_CELL_ADHESION
GOBP_CELL_DIVISION
F
Multi-Bubble Plot of GSEA Results
Runx1+ SCT MERGED (90G)
Runx1+ LOG MERGED (209G)
-4 -2 0 2 4 6 8
Normalized Enrichment Score (NES)
Comparison
4W WT vs Runx1 KO C1 MaF
4W WT vs Runx1 KO C2 QuieCF
4W WT vs Runx1 KO C3 ActCF-1
4W WT vs Runx1 KO C4 ActCF-2
4W WT vs Runx1 KO C5 ActCF-3
4W WT vs Runx1 KO all CF
-log10(FDR)
1
2
3
4
G
GOBP_POSITIVE_REGULATION_OF_PROTEIN_MODIFICATION_PROCESS
GOBP_REGULATION_OF_CELL_ADHESION
GOBP_HEART_MORPHOGENESIS
GOBP_NITROGEN_COMPOUND_TRANSPORT
GOBP_CHEMICAL_HOMEOSTASIS
GOBP_CIRCULATORY_SYSTEM_DEVELOPMENT
Runx1+ SCT merged
GOBP_BONE_DEVELOPMENT
GOBP_EMBRYO_DEVELOPMENT
GOBP_NEGATIVE_REGULATION_OF_DEVELOPMENTAL_PROCESS
GOBP_OSTEOBLAST_DIFFERENTIATION
H
3D_C7 ProCF
3D_C1 CMF-1
3D_C4 CMF-2
3D_C3 ActCF-1
3D_C2 QuieCF
3D_C5 ActCF-2
3D_C6 ActCF-3
7D_C7 ProCMF
7D_C4 CMF-2
7D_C3 CMF-1
7D_C6 ActCF-3
7D_C1 ActCF-1
7D_C2 QuieCF
7D_C5 ActCF-2
4W_C1 MaF
4W_C4 ActCF-2
4W_C5 ActCF-3
4W_C3 ActCF-1
4W_C2 QuieCF
I
3D_C1 CMF-1
3D_C4 CMF-2
3D_C3 ActCF-1
3D_C2 QuieCF
3D_C5 ActCF-2
3D_C6 ActCF-3
7D_C3 CMF-1
7D_C4 CMF-2
7D_C1 ActCF-1
7D_C6 ActCF-3
7D_C2 QuieCF
7D_C5 ActCF-2
4W_C1 MaF
4W_C4 ActCF-2
4W_C3 ActCF-1
4W_C2 QuieCF

**Figure S4. *Runx1* promotes CF differentiation after MI.** (A) A dot plot shows the expression of representative marker genes used for cell type annotation in cells from WT and *Runx1* KO hearts at 3D, 7D, and 4W post-MI. (B) A dot plot displays expression of selected genes in CF clusters at 3D post-MI. (C) A graph shows GOBP term clusters enriched in WT versus *Runx1* KO CMFs at 3D post-MI. (D) A multi-bubble plot shows the differential enrichment of Runx1 regulons between WT and *Runx1* KO CFs at 3D post-MI. (E) A graph shows overlap between Runx1 regulon (Runx1+ SCT merged) and core enrichment genes in GOBP terms enriched in WT versus *Runx1* KO CMF at 3D post-MI. (F) A graph shows the differential enrichment of Runx1 regulons between WT and *Runx1* KO subtypes identified at 4W post-MI. (G) A graph shows the overlap between Runx1 regulon (Runx1+ SCT merged) and core enrichment genes in GOBP terms enriched in WT versus *Runx1* KO C4 ActCF-2 at 4W post-MI. (H-I) Sankey diagrams depict predicted differentiation trajectories of WT (H) and *Runx1* KO (I) CFs across 3D, 7D, and 4W post-MI.

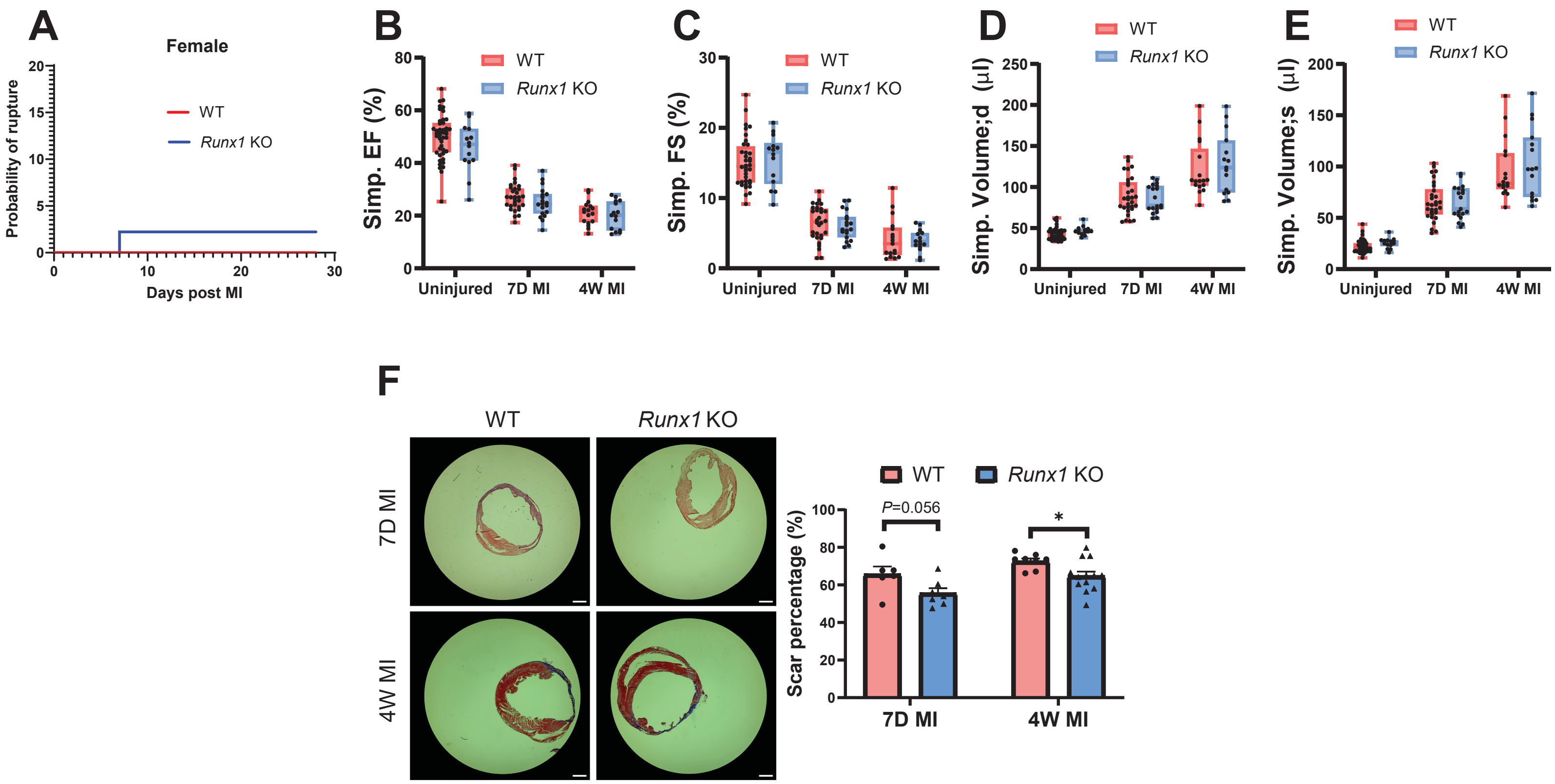
A
Female
WT
Runx1 KO
Probability of rupture
Days post MI
B
Simp. EF (%)
C
Simp. FS (%)
D
Simp. Volume;d (µl)
E
Simp. Volume;s (µl)
Uninjured
7D MI
4W MI
F
WT
Runx1 KO
7D MI
4W MI
Scar percentage (%)
P=0.056
*

**Figure S5. CF-specific *Runx1* KO ameliorates post-MI cardiac dysfunction.** (A) Post-MI rupture curves for female WT and *Runx1* KO mice. n=72 (WT), 60 (*Runx1* KO). (B-E) Ejection fraction (B), fraction shortening (C), and diastolic (D) and systolic (E) volumes of female WT and *Runx1* KO mice measured at baseline and at 7D and 4W post-MI. n=44 (uninjured WT), 14 (uninjured *Runx1* KO), 30 (7D MI WT), 19 (7D MI *Runx1* KO), 17 (4W MI WT), 15 (4W MI *Runx1* KO). (F) Trichrome staining images show the infarcted hearts from female WT and *Runx1* KO mice at 7D and 4W post-MI with quantification of infarct size. Scale bar: 1 mm. *, $p < 0.05$. n=6 (WT, 7D MI), 7 (*Runx1* KO, 7D MI), 8 (WT, 4W MI), 13 (*Runx1* KO, 4W MI).

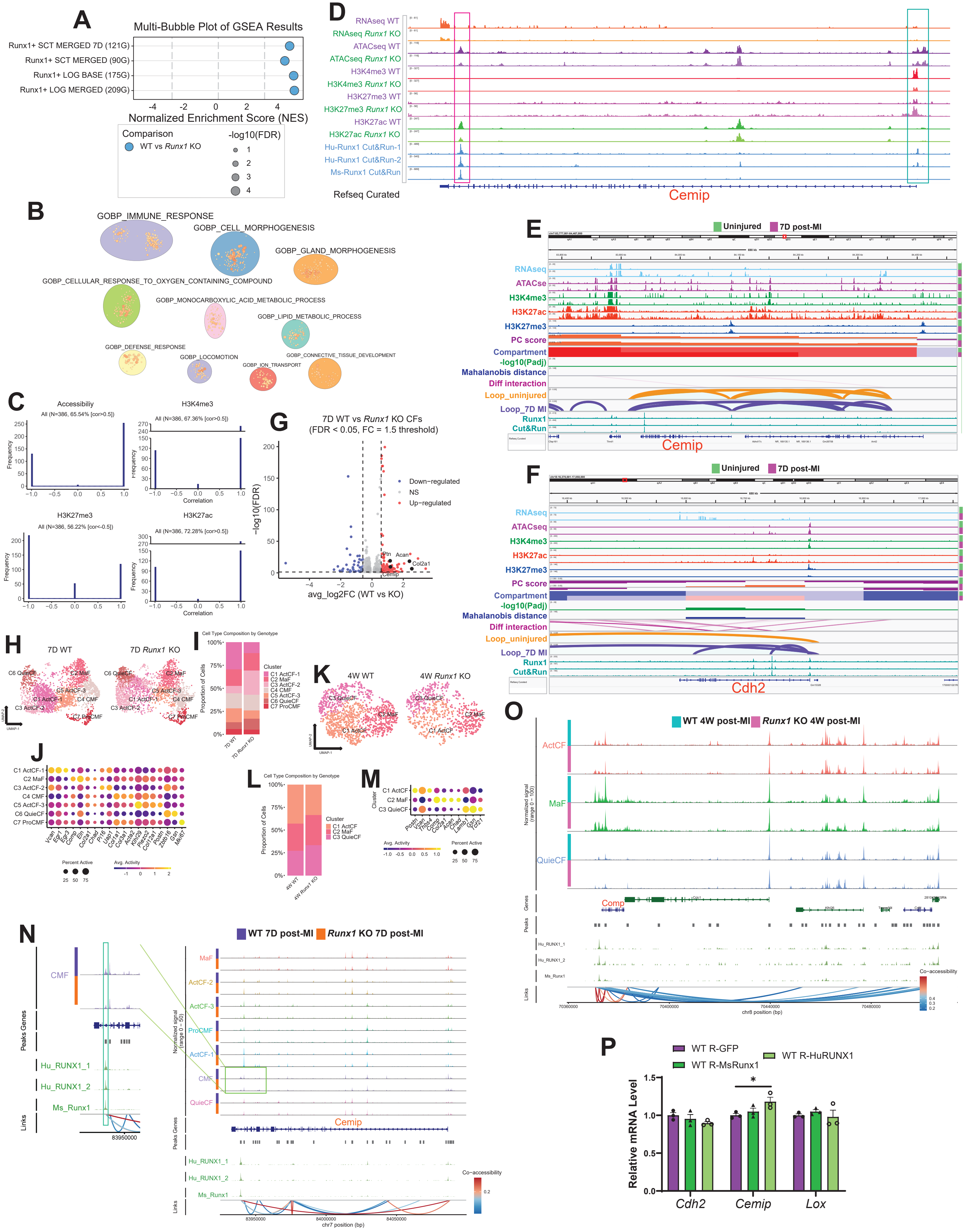
A
Multi-Bubble Plot of GSEA Results
Runx1+ SCT MERGED 7D (121G)
Runx1+ SCT MERGED (90G)
Runx1+ LOG BASE (175G)
Runx1+ LOG MERGED (209G)
Normalized Enrichment Score (NES)
Comparison
WT vs Runx1 KO
-log10(FDR)
D
RNAseq WT
RNAseq Runx1 KO
ATACseq WT
ATACseq Runx1 KO
H3K4me3 WT
H3K4me3 Runx1 KO
H3K27me3 WT
H3K27me3 Runx1 KO
H3K27ac WT
H3K27ac Runx1 KO
Hu-Runx1 Cut&Run-1
Hu-Runx1 Cut&Run-2
Ms-Runx1 Cut&Run
Refseq Curated
Cemip
B
GOBP_IMMUNE_RESPONSE
GOBP_CELL_MORPHOGENESIS
GOBP_GLAND_MORPHOGENESIS
GOBP_CELLULAR_RESPONSE_TO_OXYGEN_CONTAINING_COMPOUND
GOBP_MONOCARBOXYLIC_ACID_METABOLIC_PROCESS
GOBP_LIPID_METABOLIC_PROCESS
GOBP_DEFENSE_RESPONSE
GOBP_LOCOMOTION
GOBP_CONNECTIVE_TISSUE_DEVELOPMENT
GOBP_ION_TRANSPORT
E
Uninjured
7D post-MI
RNAseq
ATACse
H3K4me3
H3K27ac
H3K27me3
PC score
Compartment
-log10(Padj)
Mahalanobis distance
Diff interaction
Loop_uninjured
Loop_7D MI
Runx1 Cut&Run
Cemip
C
Accessibiliy
All (N=386, 65.54% [cor>0.5])
H3K4me3
All (N=386, 67.36% [cor>0.5])
H3K27me3
All (N=386, 56.22% [cor<-0.5])
H3K27ac
All (N=386, 72.28% [cor>0.5])
Frequency
Correlation
G
7D WT vs Runx1 KO CFs
(FDR < 0.05, FC = 1.5 threshold)
-log10(FDR)
avg_log2FC (WT vs KO)
Down-regulated
NS
Up-regulated
Ptn
Acan
Col2a1
Cemip
F
Uninjured
7D post-MI
RNAseq
ATACseq
H3K4me3
H3K27ac
H3K27me3
PC score
Compartment
-log10(Padj)
Mahalanobis distance
Diff interaction
Loop_uninjured
Loop_7D MI
Runx1 Cut&Run
Cdh2
H
7D WT
7D Runx1 KO
C6 QuieCF
C2 MaF
C5 ActCF-3
C1 ActCF-1
C4 CMF
C3 ActCF-2
C7 ProCMF
UMAP-1
UMAP-2
I
Cell Type Composition by Genotype
Proportion of Cells
Cluster
C1 ActCF-1
C2 MaF
C3 ActCF-2
C4 CMF
C5 ActCF-3
C6 QuieCF
C7 ProCMF
7D WT
7D Runx1 KO
K
4W WT
4W Runx1 KO
C3 QuieCF
C2 MaF
C1 ActCF
J
C1 ActCF-1
C2 MaF
C3 ActCF-2
C4 CMF
C5 ActCF-3
C6 QuieCF
C7 ProCMF
Vcan Egr1 Egr3 Comp Eln Col2a1 Chad Pi16 Uap1 Col1a1 Col3a1 Acta2 Klf29 Piezo2 Col12a1 Postn Zbtb16 Gsn Mki67
Percent Active
Avg. Activity
L
Cell Type Composition by Genotype
Proportion of Cells
Cluster
C1 ActCF
C2 MaF
C3 QuieCF
4W WT
4W Runx1 KO
M
C1 ActCF
C2 MaF
C3 QuieCF
Cluster
Postn Acan Thbs4 Comp Col2a1 Acan Chad Lama1 Gsn Tcf21
Avg. Activity
Percent Active
O
WT 4W post-MI
Runx1 KO 4W post-MI
ActCF
MaF
QuieCF
Normalized signal (range 0 - 100)
Genes
Peaks
Comp
Hu_RUNX1_1
Hu_RUNX1_2
Ms_Runx1
Links
Co-accessibility
chr8 position (bp)
N
WT 7D post-MI
Runx1 KO 7D post-MI
CMF
MaF
ActCF-2
ActCF-3
ProCMF
ActCF-1
QuieCF
Peaks Genes
Hu_RUNX1_1
Hu_RUNX1_2
Ms_Runx1
Links
Normalized signal (range 0 - 50)
Cemip
Co-accessibility
chr7 position (bp)
P
WT R-GFP
WT R-MsRunx1
WT R-HuRUNX1
Relative mRNA Level
Cdh2
Cemip
Lox

**Figure S6. Multiomic analysis identifies epigenetic regulation of CF transcriptome by *Runx1*.** (A) A multi-bubble plot shows the differential enrichment of Runx1 regulons in cultured WT versus *Runx1* KO CMFs. (B) Top 10 clusters of GOBP terms enriched in cultured WT versus *Runx1* KO CMFs. (C) Histograms show the correlations between gene expression and indicated epigenetic markers in the promoter regions for DEGs between WT and *Runx1* KO CMFs. (D) Multi-modal tracks show RNAseq and epigenetic profiles (ATACseq, histone marks) of cultured WT and *Runx1* KO CMFs, along with Runx1/RUNX1 CUT&RUN tracks at the *Cemip* locus. Cyan and pink boxes highlight Cemip promoter and enhancer, respectively. (E-F) Integrated views for QuieCFs (uninjured) and CMFs (7D post-MI) show RNAseq, ATACseq, and histone CUT&Tag (H3K4me3, H3K27ac, H3K27me3) tracks, Hi-C tracks with dcHiC-derived PC scores, compartments (dark red: sA; light red: wA; light blue: wB; dark blue: sB), compartment transition significance (-log10 Padj), Mahalanobis distance, differential interactions, Juicer-identified contacts, and Runx1/RUNX1 CUT&RUN tracks. Panels show *Cemip* (E) and *Cdh2* (F) loci. (G) A volcano plot shows DEGs between WT and *Runx1* KO CFs at 7D post-MI. (H-I) A multi-modality joint UMAP graph (H) and a bar graph (I) show different CF subtypes identified in WT and *Runx1* KO mouse hearts at 7D post-MI. (J) A dot plot shows the gene activity score of representative marker genes used for CF subtype annotation at 7D post-MI. (K-M) A multi-modality joint UMAP graph (K) and a bar graph (L) show different CF subtypes identified in WT and *Runx1* KO mouse hearts at 4W post-MI. (M) A dot plot shows the gene activity score of representative marker genes used for CF subtype annotation at 4W post-MI. (N-O) Graphs illustrate co-accessibility between regions around the *Cemip* (N) and *Comp* (O) loci identified by snMultiome analysis of WT and *Runx1* KO CFs at 7D (N) and 4W (O) post-MI. The inset in (N) highlights a Runx1-targeted intragenic enhancer located in the *Cemip* gene. (P) A bar graph shows the effects of overexpressing mouse *Runx1* (MsRunx1) and human *RUNX1* (HuRUNX1) OE on the expression of indicated Runx1 target genes in CFs.

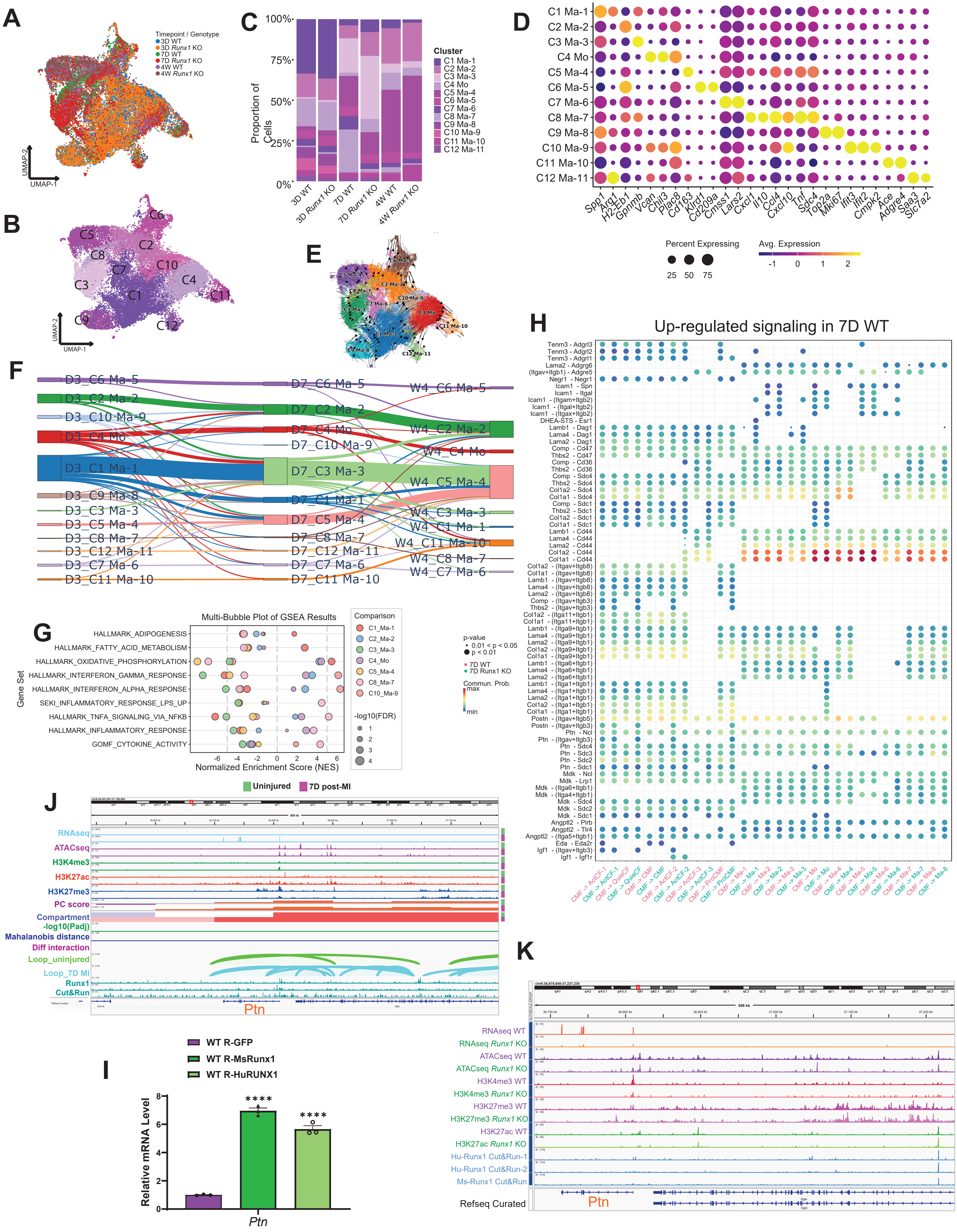

A
Timepoint / Genotype
3D WT
3D Runx1 KO
7D WT
7D Runx1 KO
4W WT
4W Runx1 KO
UMAP-2
UMAP-1
B
C1
C2
C3
C4
C5
C6
C7
C8
C9
C10
C11
C12
C
Proportion of Cells
100%
75%
50%
25%
0%
3D WT
3D Runx1 KO
7D WT
7D Runx1 KO
4W WT
4W Runx1 KO
Cluster
C1 Ma-1
C2 Ma-2
C3 Ma-3
C4 Mo
C5 Ma-4
C6 Ma-5
C7 Ma-6
C8 Ma-7
C9 Ma-8
C10 Ma-9
C11 Ma-10
C12 Ma-11
D
Spp1
Arg1
H2-Eb1
Gpnmb
Vcan
Chil3
Plac8
Cd163
Klrd1
Cd209a
Cmss1
Lars2
Cxcl1
Il10
Ccl4
Cxcl10
Tnf
Sdc4
Top2a
Mki67
Ifit3
Ifit2
Cmpk2
Ace
Adgre4
Saa3
Slc7a2
Percent Expressing
25
50
75
Avg. Expression
-1
0
1
2
E
C6 Ma-5
C5 Ma-4
C2 Ma-2
C8 Ma-7
C10 Ma-9
C3 Ma-3
C7 Ma-6
C4 Mo
C11 Ma-10
C1 Ma-1
C9 Ma-8
C12 Ma-11
F
D3_C6 Ma-5
D3_C2 Ma-2
D3_C10 Ma-9
D3_C4 Mo
D3_C1 Ma-1
D3_C9 Ma-8
D3_C3 Ma-3
D3_C5 Ma-4
D3_C8 Ma-7
D3_C12 Ma-11
D3_C7 Ma-6
D3_C11 Ma-10
D7_C6 Ma-5
D7_C2 Ma-2
D7_C4 Mo
D7_C10 Ma-9
D7_C3 Ma-3
D7_C1 Ma-1
D7_C5 Ma-4
D7_C8 Ma-7
D7_C12 Ma-11
D7_C7 Ma-6
D7_C11 Ma-10
W4_C6 Ma-5
W4_C2 Ma-2
W4_C4 Mo
W4_C5 Ma-4
W4_C3 Ma-3
W4_C1 Ma-1
W4_C11 Ma-10
W4_C8 Ma-7
W4_C7 Ma-6
G
Multi-Bubble Plot of GSEA Results
Gene Set
HALLMARK_ADIPOGENESIS
HALLMARK_FATTY_ACID_METABOLISM
HALLMARK_OXIDATIVE_PHOSPHORYLATION
HALLMARK_INTERFERON_GAMMA_RESPONSE
HALLMARK_INTERFERON_ALPHA_RESPONSE
SEKI_INFLAMMATORY_RESPONSE_LPS_UP
HALLMARK_TNFA_SIGNALING_VIA_NFKB
HALLMARK_INFLAMMATORY_RESPONSE
GOMF_CYTOKINE_ACTIVITY
Normalized Enrichment Score (NES)
-6
-4
-2
0
2
4
6
Comparison
C1_Ma-1
C2_Ma-2
C3_Ma-3
C4_Mo
C5_Ma-4
C8_Ma-7
C10_Ma-9
-log10(FDR)
1
2
3
4
H
Up-regulated signaling in 7D WT
Tenm3 - Adgrl3
Tenm3 - Adgrl2
Tenm3 - Adgrl1
Lama2 - Adgrg6
(Itgav+Itgb1) - Adgre5
Negr1 - Negr1
Icam1 - Spn
Icam1 - Itgal
Icam1 - (Itgam+Itgb2)
Icam1 - (Itgal+Itgb2)
Icam1 - (Itgax+Itgb2)
DHEA-STS - Esr1
Lamb1 - Dag1
Lama4 - Dag1
Lama2 - Dag1
Comp - Cd47
Thbs2 - Cd47
Comp - Cd36
Thbs2 - Cd36
Comp - Sdc4
Thbs2 - Sdc4
Col1a2 - Sdc4
Col1a1 - Sdc4
Comp - Sdc1
Thbs2 - Sdc1
Col1a2 - Sdc1
Col1a1 - Sdc1
Lamb1 - Cd44
Lama4 - Cd44
Lama2 - Cd44
Col1a2 - Cd44
Col1a1 - Cd44
Col1a2 - (Itgav+Itgb8)
Col1a1 - (Itgav+Itgb8)
Lamb1 - (Itgav+Itgb8)
Lama4 - (Itgav+Itgb8)
Lama2 - (Itgav+Itgb8)
Comp - (Itgav+Itgb3)
Thbs2 - (Itgav+Itgb3)
Col1a2 - (Itga11+Itgb1)
Col1a1 - (Itga11+Itgb1)
Lamb1 - (Itga9+Itgb1)
Lama4 - (Itga9+Itgb1)
Lama2 - (Itga9+Itgb1)
Col1a2 - (Itga9+Itgb1)
Col1a1 - (Itga9+Itgb1)
Lamb1 - (Itga6+Itgb1)
Lama4 - (Itga6+Itgb1)
Lama2 - (Itga6+Itgb1)
Lamb1 - (Itga1+Itgb1)
Lama4 - (Itga1+Itgb1)
Lama2 - (Itga1+Itgb1)
Col1a2 - (Itga1+Itgb1)
Col1a1 - (Itga1+Itgb1)
Postn - (Itgav+Itgb5)
Postn - (Itgav+Itgb3)
Ptn - Ncl
Ptn - (Itgav+Itgb3)
Ptn - Sdc4
Ptn - Sdc3
Ptn - Sdc2
Ptn - Sdc1
Mdk - Ncl
Mdk - Lrp1
Mdk - (Itga6+Itgb1)
Mdk - (Itga4+Itgb1)
Mdk - Sdc4
Mdk - Sdc2
Mdk - Sdc1
Angptl2 - Pirb
Angptl2 - Tlr4
Angptl2 - (Itga5+Itgb1)
Eda - Eda2r
Igf1 - (Itgav+Itgb3)
Igf1 - Igf1r
p-value
0.01 < p < 0.05
p < 0.01
7D WT
7D Runx1 KO
Commun. Prob.
max
min
CMF -> ActCF-1
CMF -> ActCF-1
CMF -> QuieCF
CMF -> QuieCF
CMF -> CMF
CMF -> CMF
CMF -> ActCF-2
CMF -> ActCF-2
CMF -> ActCF-3
CMF -> ActCF-3
CMF -> ProCMF
CMF -> ProCMF
CMF -> Ma-1
CMF -> Ma-1
CMF -> Ma-2
CMF -> Ma-2
CMF -> Ma-3
CMF -> Ma-3
CMF -> Mo
CMF -> Mo
CMF -> Ma-4
CMF -> Ma-4
CMF -> Ma-5
CMF -> Ma-5
CMF -> Ma-6
CMF -> Ma-6
CMF -> Ma-7
CMF -> Ma-7
CMF -> Ma-8
CMF -> Ma-8
J
Uninjured
7D post-MI
RNAseq
ATACseq
H3K4me3
H3K27ac
H3K27me3
PC score
Compartment
-log10(Padj)
Mahalanobis distance
Diff interaction
Loop_uninjured
Loop_7D MI
Runx1
Cut&Run
Ptn
K
RNAseq WT
RNAseq Runx1 KO
ATACseq WT
ATACseq Runx1 KO
H3K4me3 WT
H3K4me3 Runx1 KO
H3K27me3 WT
H3K27me3 Runx1 KO
H3K27ac WT
H3K27ac Runx1 KO
Hu-Runx1 Cut&Run-1
Hu-Runx1 Cut&Run-2
Ms-Runx1 Cut&Run
Refseq Curated
Ptn
558 kb
I
WT R-GFP
WT R-MsRunx1
WT R-HuRUNX1
Relative mRNA Level
8
6
4
2
0
****
****
Ptn

**Figure S7**. ***Runx1* regulates CF-macrophage communication after MI**. (A) A UMAP graph shows Mo/Ma identified in WT and *Runx1* KO hearts at 3D, 7D, and 4W post-MI, with cells color-coded by treatment groups. (B-C) A UMAP graph (B) and a bar graph (C) show different Mo/Ma subtypes identified from WT and *Runx1* KO hearts across 3D, 7D, and 4W post-MI. (D) A dotplot shows the expression of selected marker genes in Mo/Ma subtypes identified across 3D, 7D, and 4W post-MI. (E) Pseudotime trajectory of Mo/Ma from WT and *Runx1* KO mice across 3D, 7D, and 4W post-MI generated using CellRank2. (F) A Sankey graph generated using MOSCOT shows Mo/Ma differentiation trajectory in *Runx1* KO mice across indicated post-MI time points. (G) A multi-bubble plot shows the enrichment of selected GSEA terms in Mo/Ma clusters identified across 3D, 7D, and 4W post-MI. (H) A dotplot shows communications from CMF to CF and MoMa subtypes upregulated in WT versus *Runx1* KO hearts at 7D post-MI. (I) A bar graph shows the effects of *Runx1* OE on *Ptn* expression in CFs. (J) An integrated view for QuieCFs (uninjured) and CMFs (7D post-MI) show RNAseq, ATACseq, and histone CUT&Tag (H3K4me3, H3K27ac, H3K27me3) tracks, Hi-C tracks with dcHiC-derived PC scores, compartments (dark red: sA; light red: wA; light blue: wB; dark blue: sB), compartment transition significance (-log10 Padj), Mahalanobis distance, differential interactions, Juicer-identified contacts, and Runx1/RUNX1 CUT&RUN tracks around the *Ptn* locus. (K) Multi-modal tracks show RNAseq and epigenetic profiles (ATACseq, histone marks) of cultured WT and *Runx1* KO CMFs, along with Runx1/RUNX1 CUT&RUN tracks at the *Ptn* locus.

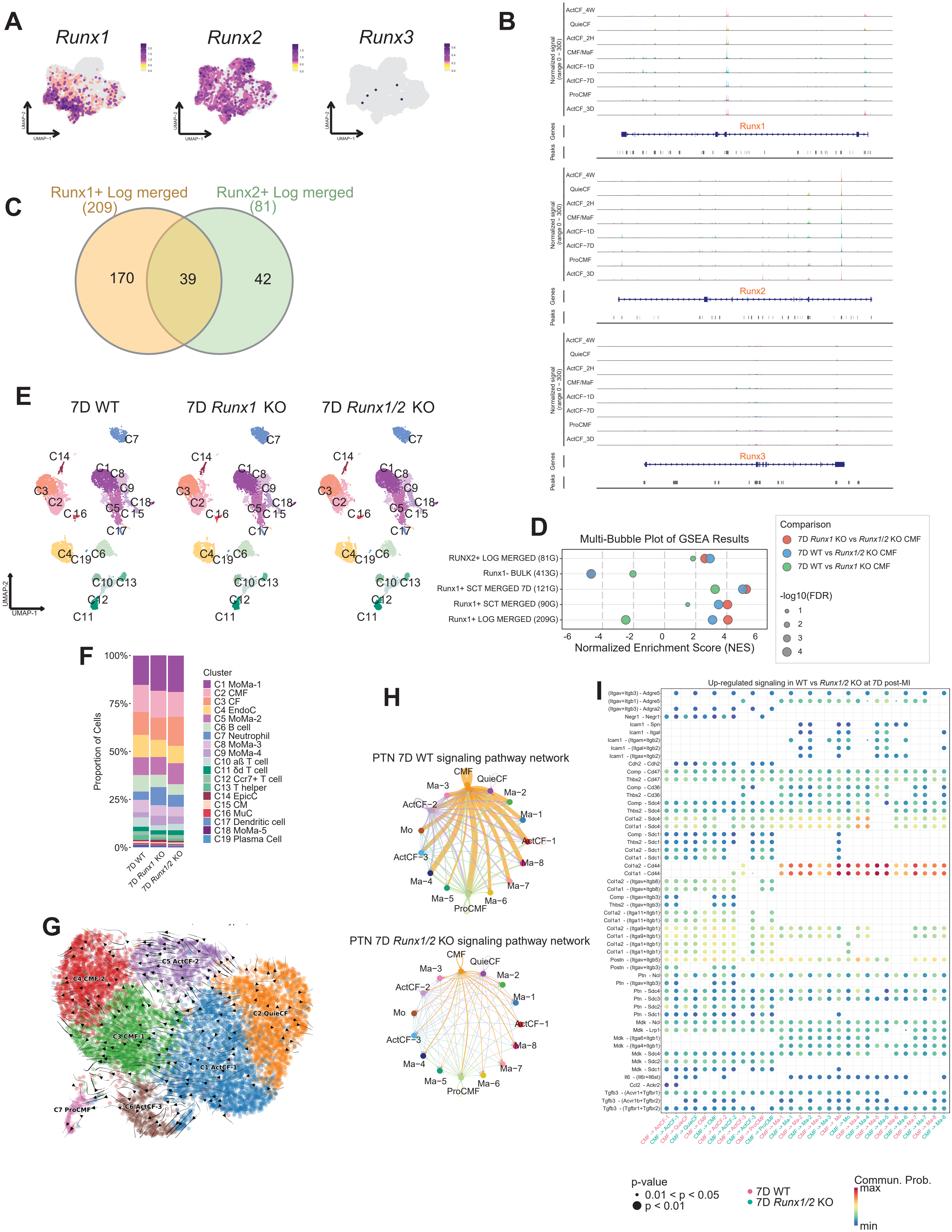

A
Runx1
Runx2
Runx3
UMAP-1
UMAP-2
B
ActCF_4W
QuieCF
ActCF_2H
CMF/MaF
ActCF−1D
ActCF−7D
ProCMF
ActCF_3D
Normalized signal (range 0 − 300)
Genes
Peaks
Runx1
Runx2
Runx3
C
Runx1+ Log merged (209)
Runx2+ Log merged (81)
170
39
42
D
Multi-Bubble Plot of GSEA Results
RUNX2+ LOG MERGED (81G)
Runx1- BULK (413G)
Runx1+ SCT MERGED 7D (121G)
Runx1+ SCT MERGED (90G)
Runx1+ LOG MERGED (209G)
Normalized Enrichment Score (NES)
Comparison
7D Runx1 KO vs Runx1/2 KO CMF
7D WT vs Runx1/2 KO CMF
7D WT vs Runx1 KO CMF
-log10(FDR)
E
7D WT
7D Runx1 KO
7D Runx1/2 KO
F
Proportion of Cells
Cluster
C1 MoMa-1
C2 CMF
C3 CF
C4 EndoC
C5 MoMa-2
C6 B cell
C7 Neutrophil
C8 MoMa-3
C9 MoMa-4
C10 aß T cell
C11 δd T cell
C12 Ccr7+ T cell
C13 T helper
C14 EpicC
C15 CM
C16 MuC
C17 Dendritic cell
C18 MoMa-5
C19 Plasma Cell
7D WT
7D Runx1 KO
7D Runx1/2 KO
G
C4 CMF-2
C5 ActCF-2
C2 QuieCF
C3 CMF-1
C1 ActCF-1
C7 ProCMF
C6 ActCF-3
H
PTN 7D WT signaling pathway network
PTN 7D Runx1/2 KO signaling pathway network
CMF
QuieCF
Ma−2
Ma−1
ActCF−1
Ma−8
Ma−7
Ma−6
ProCMF
Ma−5
Ma−4
ActCF−3
Mo
ActCF−2
Ma−3
I
Up-regulated signaling in WT vs Runx1/2 KO at 7D post-MI
p-value
0.01 < p < 0.05
p < 0.01
7D WT
7D Runx1/2 KO
Commun. Prob.
max
min

**Figure S8. Combined *Runx1* and *Runx2* KO enhances the effects of *Runx1* deletion in CFs.** (A) Feature plots show *Runx1*, *Runx2*, and *Runx3* expression in CF clusters identified from WT mice without injury and at different time points post-MI. (B) ATACseq tracks illustrate chromatin accessibility at *Runx1*, *Runx2*, and *Runx3* loci in CFs from WT hearts without injury and at different time points post-MI. (C) A Venn diagram shows overlap between Runx1 and Runx2 target genes. (D) A graph shows differential enrichment of Runx1 and Runx2 regulons among WT, *Runx1* KO, and *Runx1/2* KO CMFs. (E-F) UMAP graphs (E) and a bar graph (F) show cell types identified in WT, *Runx1* KO, and *Runx1/2* KO hearts at 7D post-MI. (G) Pseudotime trajectory of WT, *Runx1* KO, and *Runx1/2* KO CF at 7D post-MI. (H) Circle plots show Ptn-mediated signaling between CF and Mo/Ma clusters in WT and *Runx1/2* KO hearts at 7D post-MI. (I) A dot plot shows communication from CMF to CF and Mo/Ma subtypes upregulated in WT versus *Runx1/2* KO hearts at 7D post-MI.
